# CuticleTrace: A toolkit for capturing cell outlines of leaf cuticle with implications for paleoecology and paleoclimatology

**DOI:** 10.1101/2023.07.23.550217

**Authors:** Benjamin A. Lloyd, Richard S. Barclay, Regan E. Dunn, Ellen D. Currano, Ayuni I. Mohamaad, Kymbre Skersies, Surangi W. Punyasena

## Abstract

**Premise:** Leaf epidermal cell morphology is closely tied to plants’ evolutionary histories and growth environments, and is therefore of interest to many plant biologists. However, cell measurement can be time-consuming and restrictive with current methods. CuticleTrace is a suite of FIJI and R-based functions that streamlines and automates the segmentation and measurement of epidermal pavement cells across a wide range of cell morphologies and image qualities.

**Methods and Results:** We evaluated CuticleTrace-generated measurements against those from alternate automated methods and expert and undergraduate hand-tracings across a taxonomically diverse 50-image dataset of variable image qualities. We observed ∼93% statistical agreement between CuticleTrace and expert hand-traced measurements, outperforming alternate methods.

**Conclusions:** CuticleTrace is broadly applicable, modular, and customizable, and integrates data visualization and cell shape measurement with image segmentation, lowering the barrier to high-throughput studies of epidermal morphology by vastly decreasing the labor investment required to generate high-quality cell shape datasets.

## INTRODUCTION

The accurate, consistent, and efficient characterization of the leaf epidermis is key to advancing our understanding of the relationships of plants to their environments and evolutionary histories. The morphology and arrangement of leaf epidermal cells varies significantly across different environmental conditions and taxonomic groups (Kürschner, 1997; Royer, 2001; Vőfély et al., 2019). These relationships have been used to constrain paleoenvironmental conditions such as atmospheric CO_2_ concentration (McElwain and Chaloner, 1996; Royer, 2001) and canopy structure (Dunn et al., 2015; Bush et al., 2017; Milligan et al., 2021), and to enhance taxonomic resolution of paleofloras (e.g. Strömberg, 2011). In crop science, phenomics links high-throughput phenotyping of leaf epidermal characters (cell size, shape, number, etc.) with genomics to breed plants with idealized physiological traits (Zhao et al., 2019).

Efforts to automate leaf epidermal measurement have met differing levels of success. Stomata can be efficiently identified from microscope images (e.g., Fetter et al., 2019; Li et al., 2022), but the accurate characterization of epidermal pavement cell morphologies remains limited to specific taxa or imaging techniques (Möller et al., 2017; Li et al., 2022). As a result, hand-tracing has remained the most viable option for characterizing epidermal pavement cell morphology in most cases (e.g., Vőfély et al., 2019; Brown and Jordan, 2023). However, hand-tracing is a slow and tedious process, with cell-outline quality highly dependent upon both the tools used for tracing (e.g., mouse, stylus, tablet, desktop computer) and the experience level and conscientiousness of the tracer. The high labor requirement of hand tracing—and the non-standardized measurements that result—imposes a barrier to the use of cell shape in multiple disciplines. Regardless, hand-traced datasets have been part of high-impact studies studying cell shape, growth environment, and evolutionary history (Dunn et al., 2015; Bush et al., 2017; Vőfély et al., 2019). These studies demonstrate how cell shape datasets enable significant developments in paleobotany and evolutionary biology.

We developed the CuticleTrace toolkit to streamline and automate the tracing and measurement of epidermal pavement cells across a wide variety of taxa, image qualities, and image preparations. Our easily fine-tuned open-source workflow uses the freeware software FIJI (Schindelin et al., 2012) and R Statistical Software (R) (R Core Team, 2022). CuticleTrace generates large cell-shape datasets from images of leaf cuticle or epidermis. Cell-tracing is fast, consistent, and reproduces expert-level measurements.

## METHODS AND RESULTS

### CuticleTrace description

The CuticleTrace workflow consists of four steps. First, users determine appropriate batch-processing inputs with the “Single Image Processor” FIJI macro (Fig. 1b). Second, users batch-process images and measure cells with the “Batch Generate ROIs” and “Batch Measure (Different Scales)” FIJI macros, which generate (1) thresholded and skeletonized binary images (Fig. 1f-g), (2) sets of files recording individual cell shapes known as “regions of interest” (ROIs; Fig. 1i), and (3) shape parameter measurements associated with each ROI (Fig. 1j). Third, resulting cell measurements are filtered with median statistics by the “CuticleTrace Data Filtration” R notebook (Fig. 1k). Last, users can visualize the effects of data filtration with the “Batch Overlay” FIJI macro (Fig. 1j).

**Fig. 1.**
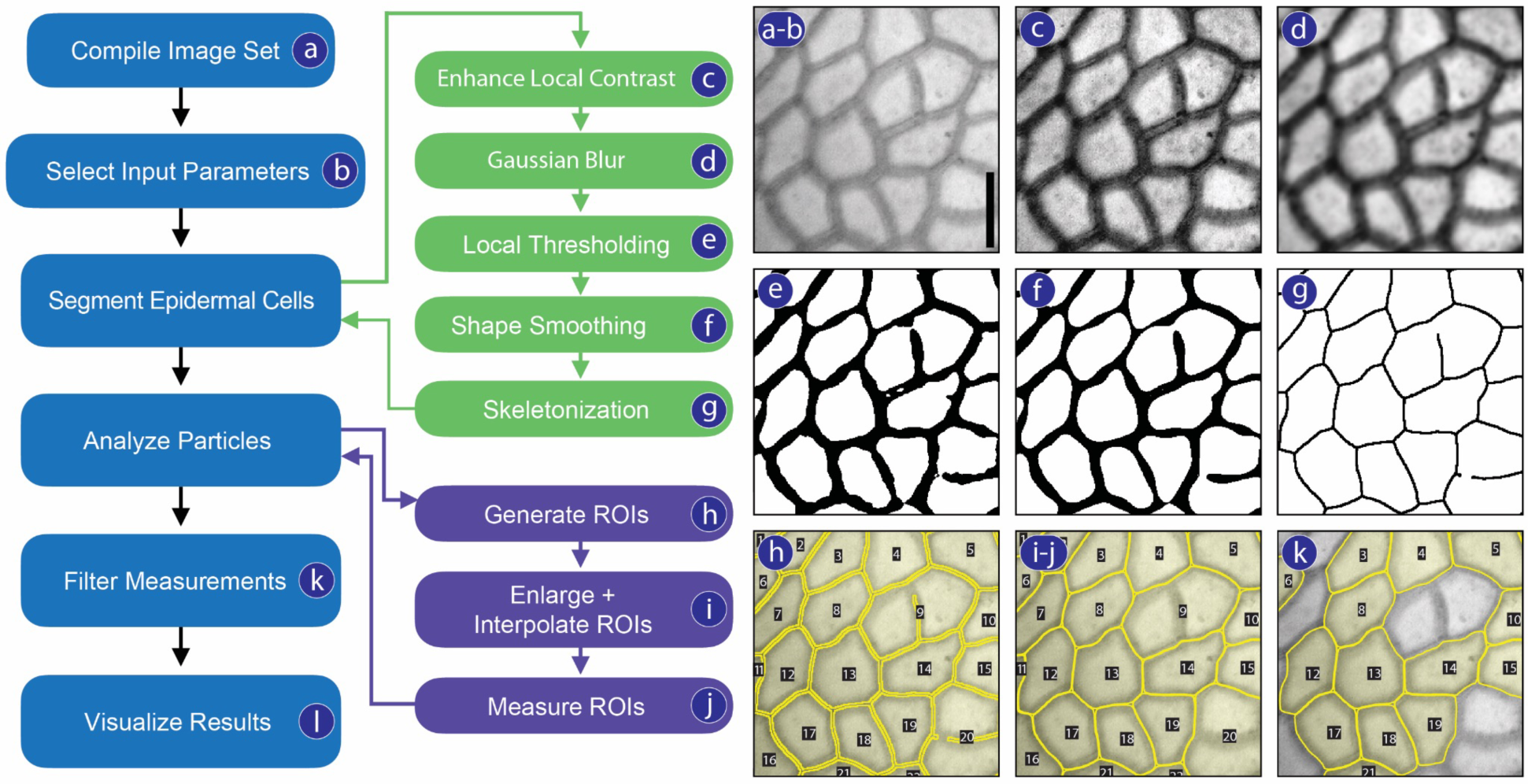
Processing workflow for the CuticleTrace toolkit (left), illustrated on an image of *Castanea pumila* (Fagaceae, FLMNH00115). Scale bar is 10 µm. Processing starts by **(a)** compiling an image set and **(b)** selecting input parameters. This is followed by epidermal cell segmentation, consisting of **(c)** local contrast enhancement, **(d)** Gaussian blur, **(e)** local thresholding, **(f)** shape smoothing, and **(g)** skeletonization. Particles are analyzed in FIJI, by **(h)** generating ROIs, **(i)** enlarging and interpolating ROIs, and **(j)** measuring ROIs. ROI measurements are **(k)** filtered with median statistics, and **(l)** visualized overlaying unprocessed images.

An illustrated manual for installation and use of FIJI and R tools is available in the supporting information (**Appendix S1**). A video tutorial is available online and all software is documented on the CuticleTrace GitHub repository (https://github.com/benjlloyd/CuticleTrace).

#### Determination of batch processing inputs

Both the “Single Image Processor” and “Batch Generate ROIs” macros apply seven user inputs to images. “Single Image Processor” works with an open image, while “Batch Generate ROIs” processes all images in an image set. Each user input sets the parameters for different image processing operations. Before batch-processing, users must first determine the appropriate input parameters on a small subset of images using the “Single Image Processor” macro. The selection of appropriate input parameters is extremely important to ensure accurate results (Fig. 2).

**Fig. 2.**
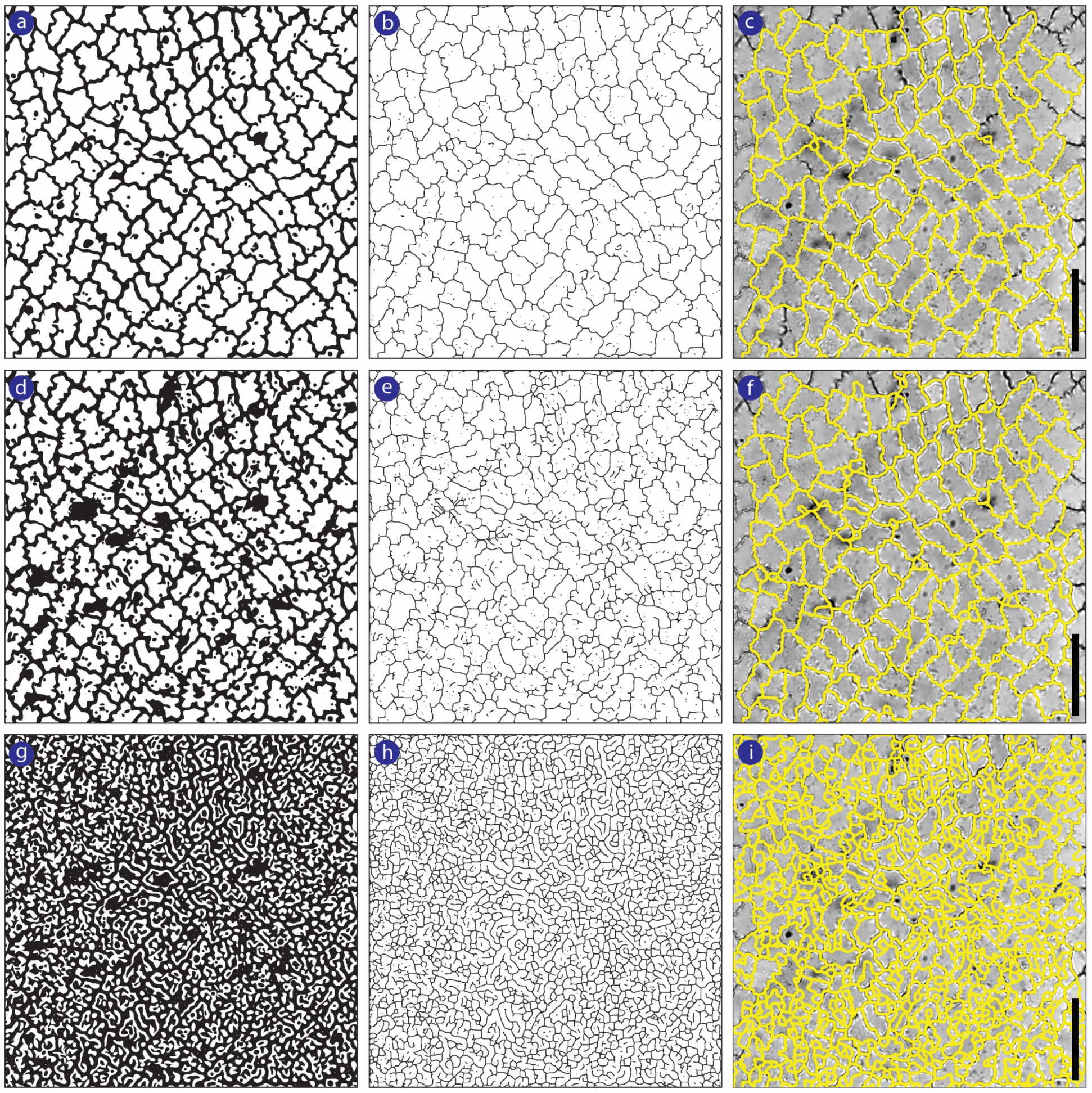
The effect of thresholding **(a, d, g)** on skeletonization **(b, e, h)** and unfiltered ROI sets **(c, f, i)**, demonstrated on *Talisia princeps* - Sapindaceae, FLMNH00443. The image is thresholded by **(a-c)** Sauvola local thresholding (ideal), **(d-f)** Bernsen local thresholding (satisfactory), and **(g-i)** NiBlack local thresholding (ineffective). Scale bars are 50µm.

For both macros, users select:

1. Whether cell walls are dark on a light background, or light on a dark background
2. The standard deviation of Gaussian blur (Fig. 1c)
3. The automated local thresholding algorithm (Fig. 1d, Fig. 2)
4. The initial local thresholding radius (Fig. 1d)
5. The range of sizes (in pixels squared) of all cells expected in the image set (Fig. 1d)
6. The percentage of Fourier descriptors to retain during shape smoothing (Fig. 1e)
7. The scale (in pixels/unit) of all images in the image set (Fig. 1i)

#### Image processing and measurement

The “Batch Generate ROIs” macro outputs thresholded and skeletonized binary images, and unfiltered (enhanced and interpolated) ROI sets of all images in a directory (Fig. 1c-i). Users may also elect to output measurements of ROI shape parameters if all images have the same scale, or use “Batch Measure (Different Scales)” to measure ROIs of images with differing scales (Fig. 1j, Table 1). These unfiltered ROI sets and measurement files may then be brought directly into the “CuticleTrace Data Filtration” R Notebook for statistical filtration.

**Table 1.**
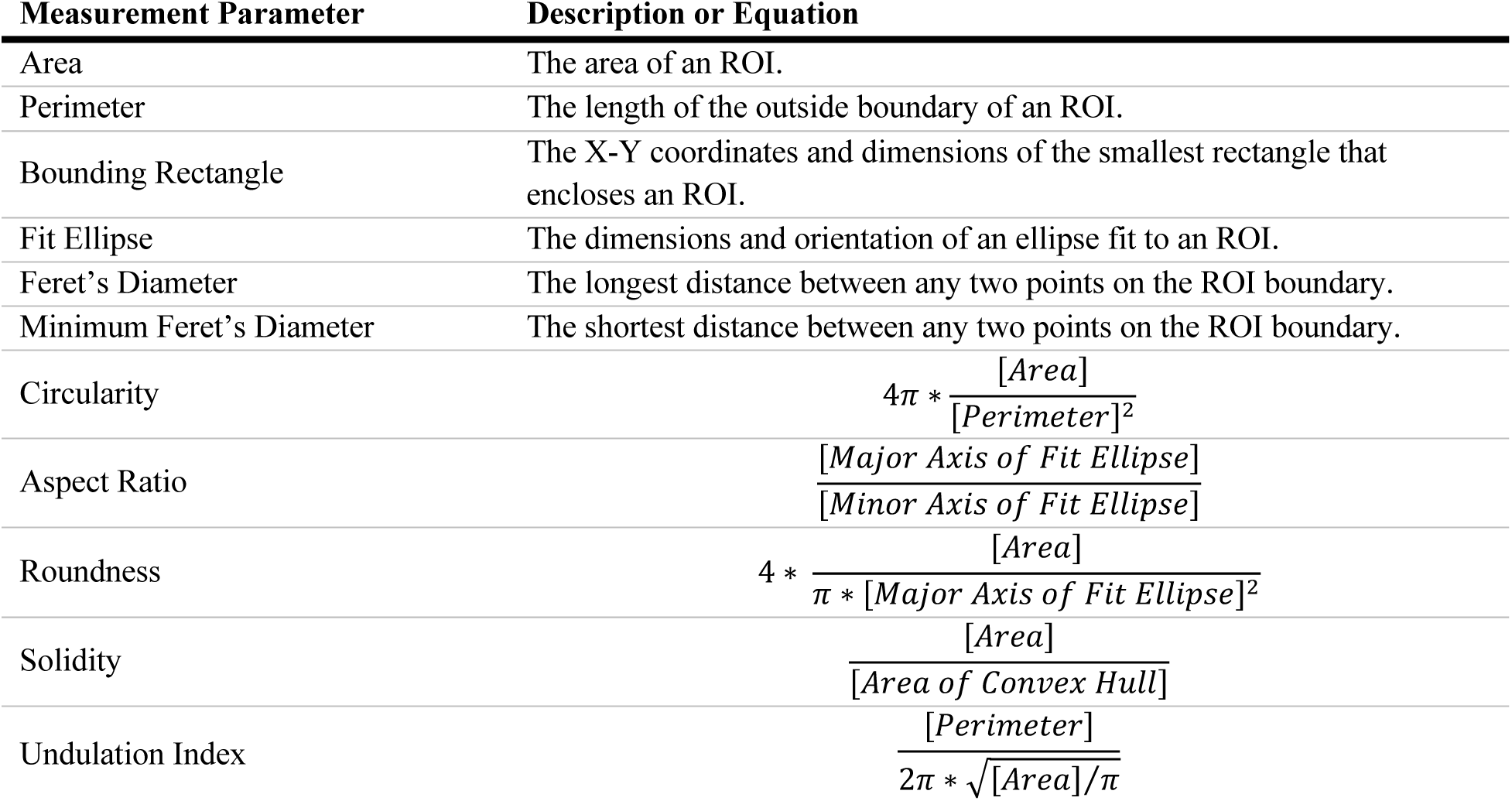
Descriptions or equations of all shape parameters measured by CuticleTrace.

#### Data filtration

The ROIs produced by the CuticleTrace FIJI analysis pipeline will inevitably include partial cells, multiple cells, vein cells, or non-cuticle image artifacts (e.g., slide background, debris). The “CuticleTrace Data Filtration” R notebook removes these erroneous data points by excluding ROIs with measurements outside one or two median absolute deviations (MADs) from each image’s median area, perimeter, circularity, aspect ratio, roundness, and solidity (Table 1). The notebook then creates new ROI sets of the remaining ROIs post-filtering (Fig. 3). We use the median as our reference point, as unfiltered measurements are non-normally distributed (Fig. 4). This approach retains ROIs that capture the central tendency of each specimen’s epidermal pavement cell morphology.

**Fig. 3.**
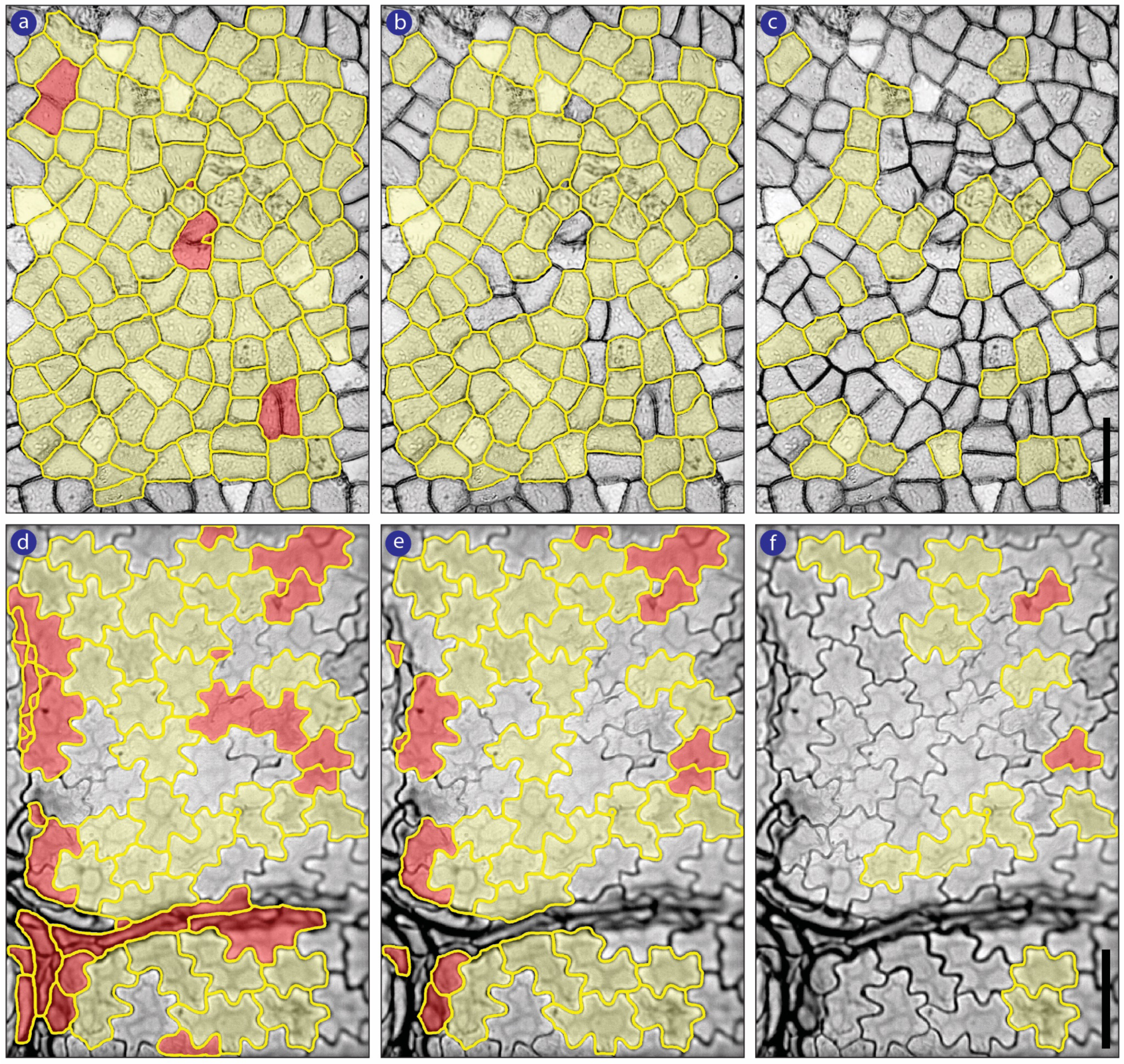
Removal of unwanted ROIs by filtering with median statistics. **(a-c)** *Ocotea tarapotana -* Lauraceae, FLMNH00185; **(d-f)** *Acer skutchii -* Aceraceae, FLMNH00714. ROI sets that are **(a, d)** unfiltered, **(b, e)** ±2MAD-filtered, and **(c, f)** ±1MAD-filtered. Yellow outlines represent ROIs retained at each filtering level. Yellow-shaded ROIs accurately characterize epidermal pavement cells. Red-shaded ROIs do not accurately characterize pavement cells and are thus not wanted in the final dataset. Scale bars are 50 µm.

**Fig. 4.**
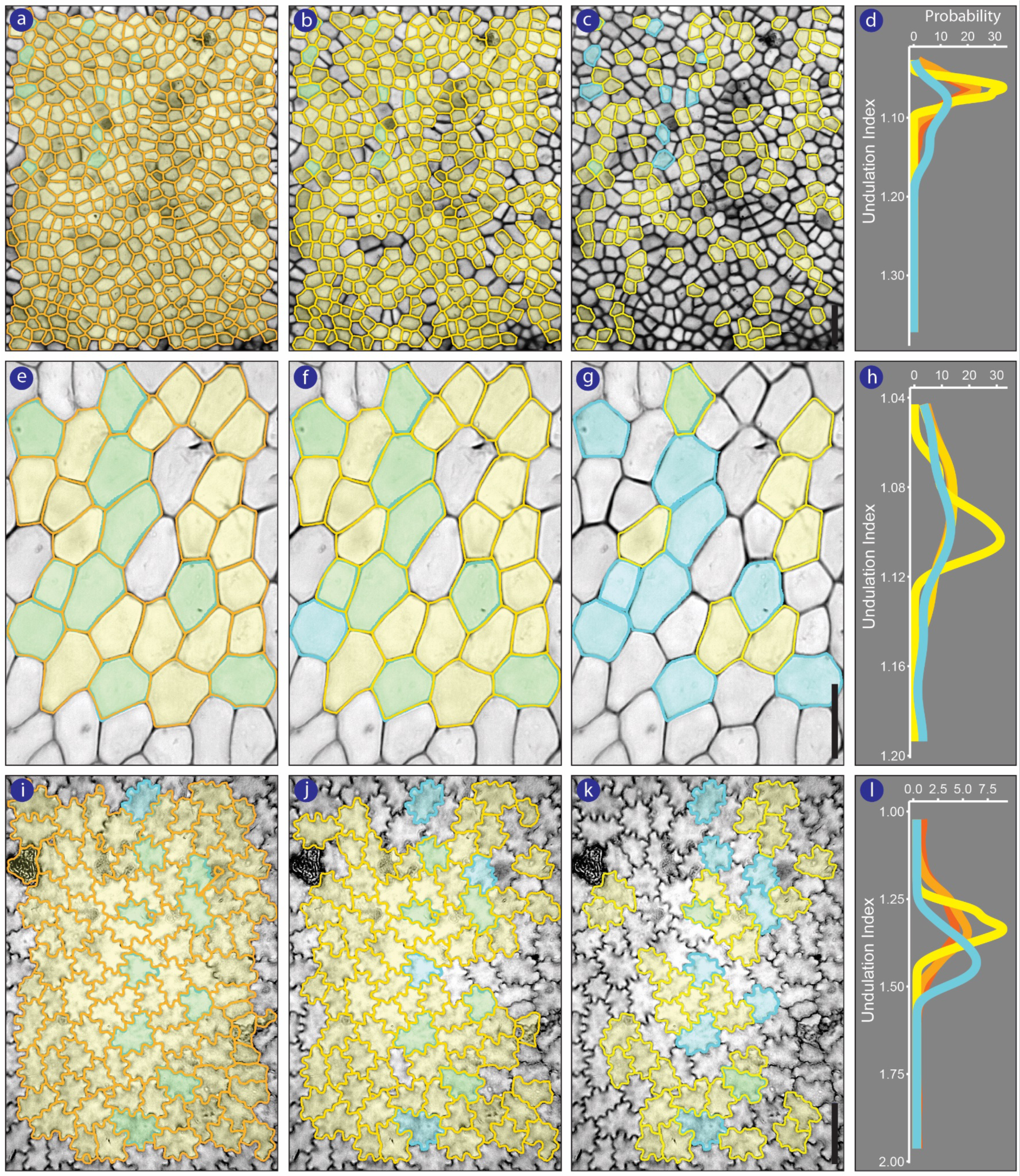
CuticleTrace-generated ROI sets (yellow) compared to expert hand drawn ROI sets (cyan), with subsequent filtering using median statistics. Cells with overlapping CuticleTrace and expert-generated ROIs appear green. Columns show ROI sets that are unfiltered **(a, e, i)**, ±2MAD **(b, f, j)**, and ±1MAD **(c, g, k)**, overlain on the source images of **(a-c)** *Nectandra oppositifolia* - Lauraceae, FLMNH00260; **(e-g)** *Guarea bijuga* - Meliaceae, FLMNH00850, and **(i-l)** *Anaxagorea petiolata* - Annonaceae, FLMNH02589. **(d, h, l)** probability density plots showing distributions of Undulation Index values across all 4 datasets for each image; hand-traced - cyan, unfiltered - dark orange, ±2MAD - light orange, ±1MAD - yellow. Scale bars are 50 µm.

#### Data visualization

The “Batch Overlay” FIJI macro allows users to visualize CuticleTrace outputs by creating images overlain with outlines of each ROI in an ROI set, formatted to the user’s preference. “Batch Overlay” may be used to overlay any batch of ROI sets on any batch of images. In our evaluation, we used “Batch Overlay” to visually check the accuracy of ROIs resulting from different input parameters (Fig. 2) and to compare unfiltered ROI sets with the filtered versions from the “CuticleTrace Data Filtration” R Notebook (Fig. 3, 4).

### CuticleTrace evaluation

We evaluated CuticleTrace across a range of image resolutions, magnifications, and qualities, using an image set of 50 vouchered herbarium specimens from the Cuticle Database (<u>cuticledb.eesi.psu.edu</u>, Barclay et al., 2007). Images were chosen to maximize taxonomic breadth (48 species, 37 genera, 20 families) and span the range of cell shapes and sizes (**Fig. S2**).

We compared CuticleTrace to alternate measurement methods in two ways. We generated CuticleTrace measurements following the protocol outlined in Appendix S1, with user inputs specified in Table 2. First, we compared CuticleTrace measurements (at the ±1MAD and ±2MAD filtering levels) to hand-traced outlines of ten cells followed Dunn et al. (2015), traced by an expert (R.E.D) and University of Wyoming undergraduates (A.I.M, K.S.); and measurements generated by alternate automated methods (LeafNet (Li et al., 2022) and PaCeQuant (Möller et al., 2017)). PaCeQuant did not successfully segment our light-microscopy images (unsurprising, as it was developed for confocal microscopy images), and was excluded from statistical comparisons of segmentation methods.

**Table 2.**
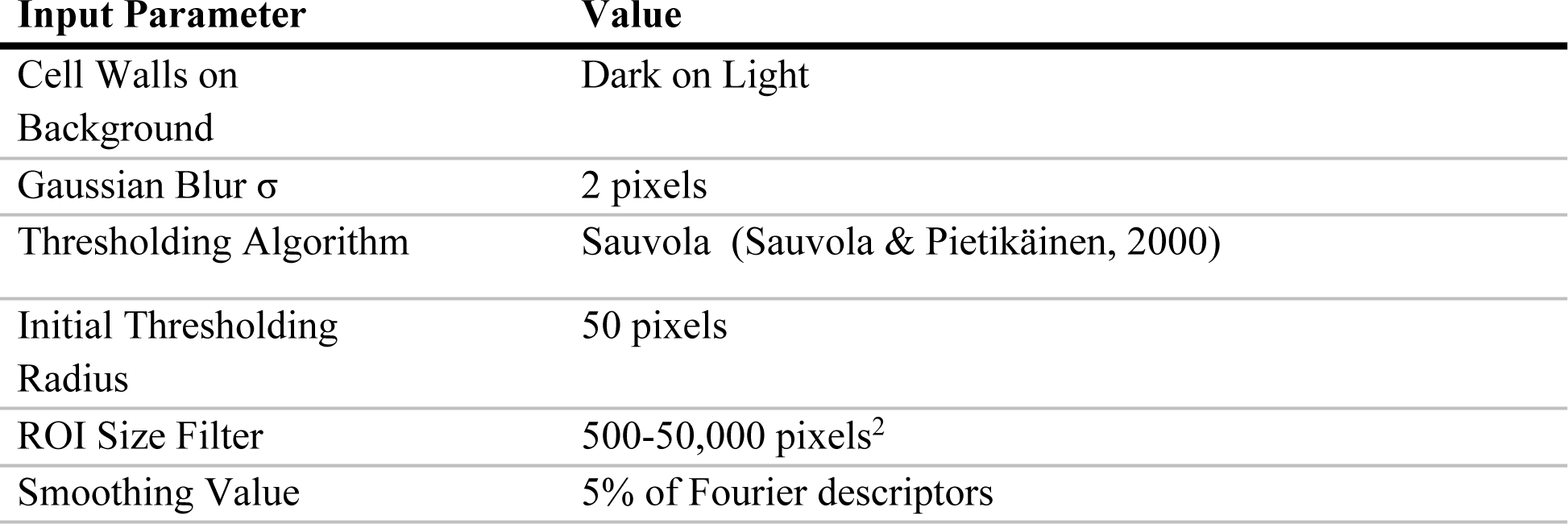
CuticleTrace batch-processing inputs for our 50-image test dataset. Input Parameter Value.

| Input Parameter | Value |
| --- | --- |
| Cell Walls on Background | Dark on Light |
| Gaussian Blur $\sigma$ | 2 pixels |
| Thresholding Algorithm | Sauvola (Sauvola & Pietikäinen, 2000) |
| Initial Thresholding Radius | 50 pixels |
| ROI Size Filter | 500-50,000 pixels<sup>2</sup> |
| Smoothing Value | 5% of Fourier descriptors |

Second, we manually reduced our CuticleTrace and expert datasets to only include the same cells (447 cells across all 50 images). We then completed a one-to-one comparison of CuticleTrace and expert measurements across all images, controlling for cell selection and sample size.

#### Whole image comparison

The expert outlines served as our evaluation benchmark. We ran a one-way Analysis of Variance (ANOVA) with post-hoc nonparametric Games-Howell analysis for each shape parameter for each image. We selected the Games-Howell test due to differing sample sizes between the four sets of measurements. We used CuticleTrace’s “Batch Overlay” macro to visually compare CuticleTrace and expert tracings (**Fig. S2**).

Our analyses showed 100% statistical agreement between CuticleTrace measurements (at both ±1MAD and ±2MAD filtering levels) and expert measurements of area, perimeter, Feret’s diameter, aspect ratio, and roundness (Fig. 5). We observed some variability in circularity, solidity and UI measurements, which was expected due to their sensitivity to small tracing differences. CuticleTrace measurements consistently aligned more closely with expert measurements than expert measurements did with student and LeafNet measurements (Table 3).

**Fig. 5.**
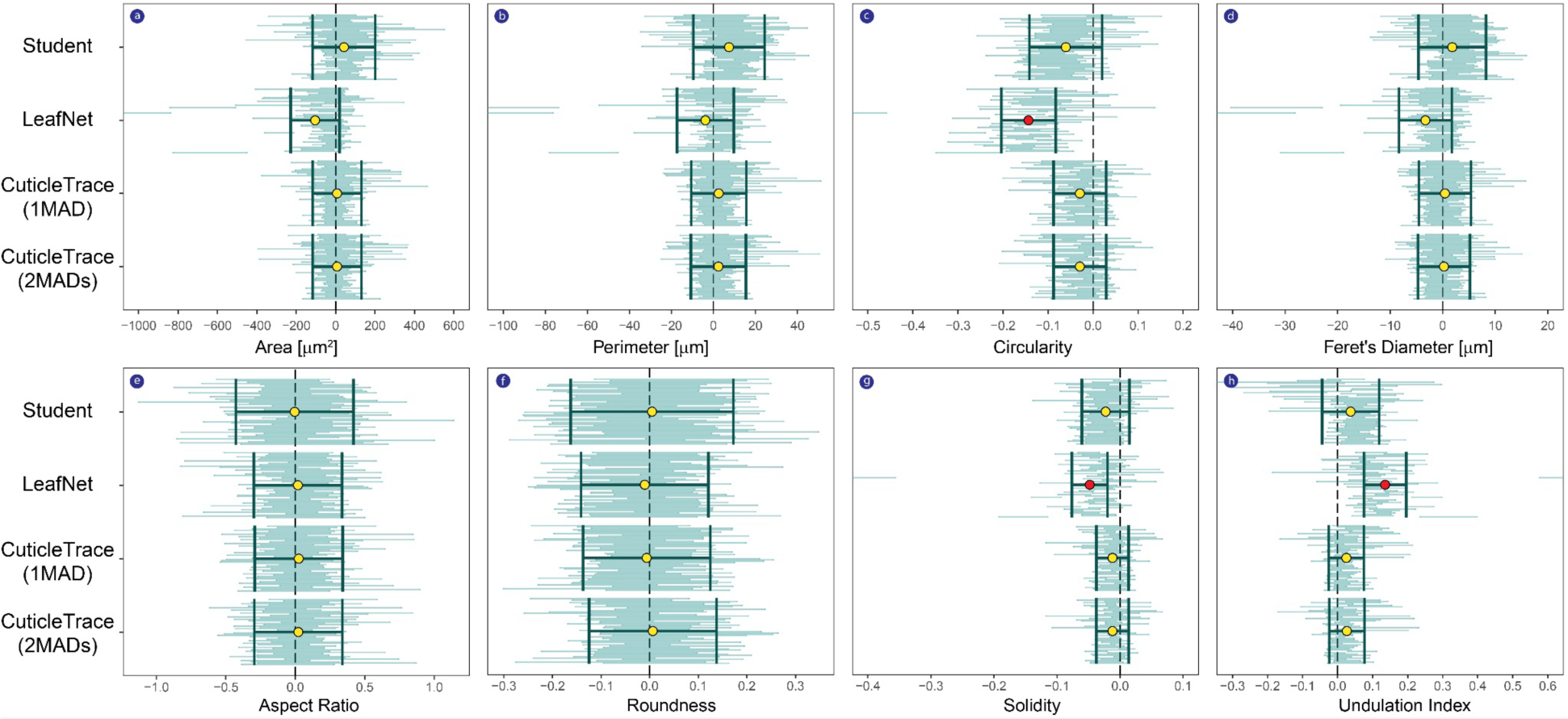
Post-hoc Games-Howell results for all 50 images, separated by shape parameter. Comparisons are all relative to the hand-drawn expert results (zero). Student data was also hand-drawn; LeafNet and CuticleTrace are automated methods. Each small horizontal bar represents the 95% confidence interval of the Games-Howell mean difference estimate applied to a single image in the comparative dataset. The image position in the stack is maintained for each shape parameter. Darker bars with center circles are the mean ±2**σ** for each of the comparative stacks of Games-Howell results. Yellow center circles are non-significant; red center circles are significantly different from the expert.

**Table 3.** Statistical evaluation of ±1MAD and ±2MAD CuticleTrace measurements, student measurements, and LeafNet measurements, in comparison to expert measurements.

| Shape<br>Parameter | Average Games-Howell p-value in comparison to<br>Expert |  |  |  |  | % of measurements statistically identical<br>to Expert (p-values >.05) |  |  |  |  |
| --- | --- | --- | --- | --- | --- | --- | --- | --- | --- | --- |
|  | CuticleTrace |  |  | Student | LeafNet | CuticleTrace |  |  | Student | LeafNet |
| | $\pm 1$MAD | $\pm 2$MAD | Cell to<br>Cell | | | $\pm 1$MAD | $\pm 2$MAD | Cell to<br>Cell | | |
| Area | 0.85 | 0.89 | 0.56 | 0.74 | 0.55 | 100 | 100 | 100 | 94 | 84 |
| Perimeter | 0.82 | 0.82 | 0.68 | 0.54 | 0.60 | 100 | 100 | 100 | 86 | 82 |
| Feret's<br>Diameter | 0.85 | 0.88 | 0.64 | 0.66 | 0.72 | 100 | 100 | 100 | 94 | 88 |
| Circularity | 0.52 | 0.53 | 0.44 | 0.31 | 0.06 | 78 | 82 | 94 | 58 | 12 |
| Aspect Ratio | 0.79 | 0.82 | 0.88 | 0.75 | 0.81 | 100 | 100 | 100 | 100 | 100 |
| Roundness | 0.78 | 0.82 | 0.88 | 0.81 | 0.78 | 100 | 100 | 100 | 100 | 100 |
| Solidity | 0.44 | 0.47 | 0.42 | 0.34 | 0.18 | 76 | 78 | 90 | 72 | 30 |
| Undulation<br>Index | 0.52 | 0.53 | 0.44 | 0.25 | 0.06 | 82 | 88 | 94 | 52 | 12 |
| Mean | 0.70 | 0.72 | 0.62 | 0.55 | 0.47 | 92.0 | 93.5 | 97.3 | 82.0 | 63.5 |

#### Cell-to-cell comparison

Our cell-to-cell comparison of CuticleTrace and expert measurements of individual cells (447 cells across all 50 images), which showed broad agreement across all shape parameters (Fig. 6). In only 11 of 400 image measurements (50 images, eight shape parameters) were values significantly different at the 2-sigma level—a 2.75% error rate. Seven of the 11 significant differences are concentrated in three images—FLMNH00178 (*Damburneya salicifolia*), FLMNH00561 (*Gleditsia triacanthos*), and FLMNH05215 (*Pouteria durlandii*) (Fig. 6). However, overall, differences between expert and CuticleTrace tracings of the same cells were small, and CuticleTrace tracings were reasonable and consistent even when they differed from expert tracings.

**Fig. 6.**
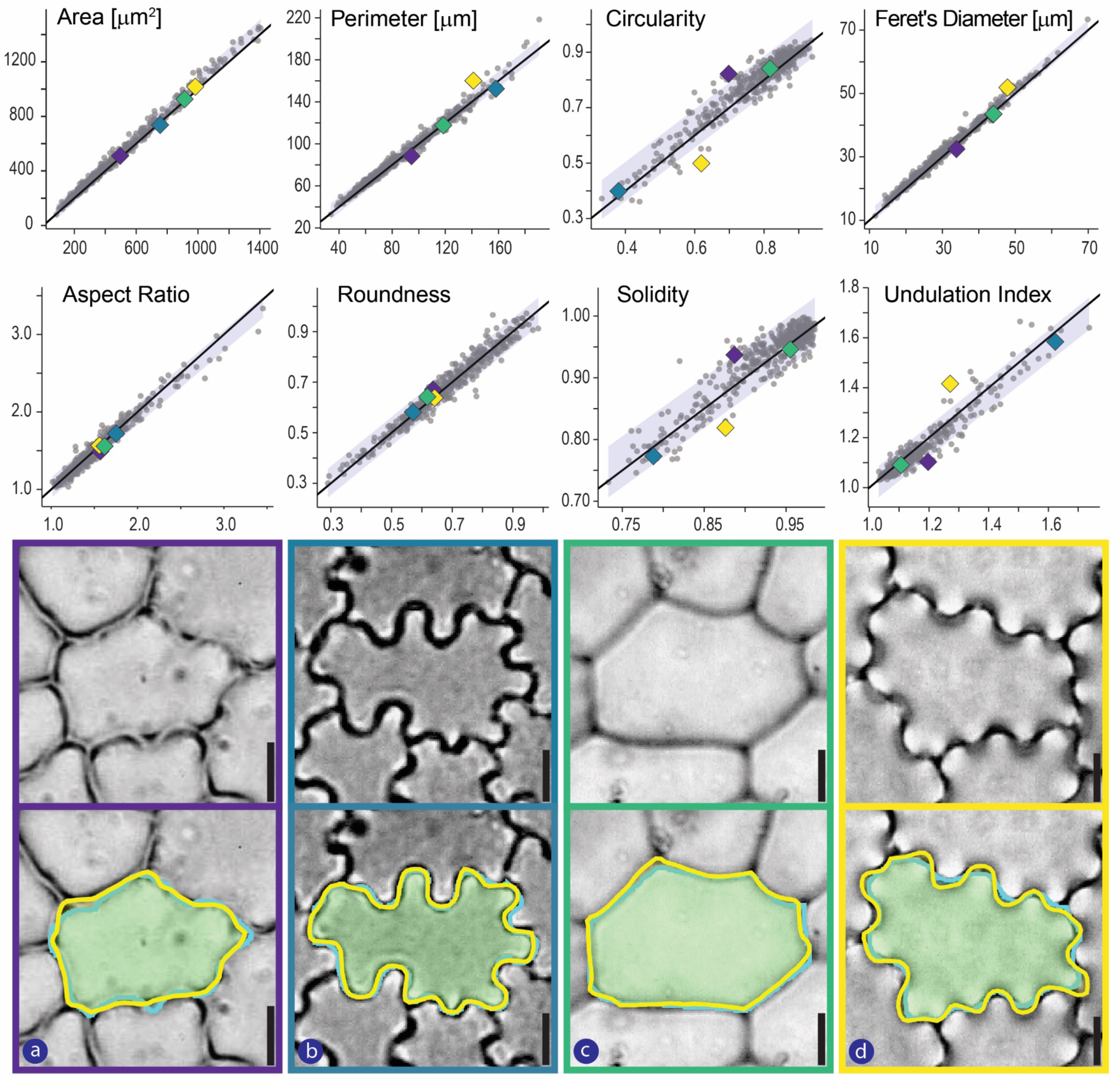
Cell-to-cell comparison of Expert and CuticleTrace measurements. Upper panel: Correlation between Expert (x-axis) and CuticleTrace (y-axis) measurements across 447 cells and all relevant shape parameters, distributed between 50 images. Black lines have a slope of one, indicating 100% correlation. Shaded areas show 95% prediction intervals. Colored symbols correlate to the four images presented in the lower panel, which provides a visual comparison of the difference between cell outlines hand-drawn by an expert (cyan line) versus the CuticleTrace automated procedure (yellow line). Lower panel: (a) *Gleditsia triacanthos* - Fabaceae, FLMNH00081; (b) *Toxicodendron striatum* - Anacardiaceae, FLMNH00510; (c) *Oreopanax capitatus* - Araliaceae, FLMNH00767; (d) *Pouteria durlandii* - Sapotaceae, FLMNH05215. Scale bars are 10 µm.

### Accuracy of CuticleTrace Measurements

Across the wide range of cell shapes and image qualities encapsulated in our test dataset (**Fig. S2**), CuticleTrace produces cell shape measurements that are statistically identical to expert measurements across all shape parameters in most instances, at both the ±1MAD and ±2MAD filtering level. The limited, often non-significant, differences between expert and CuticleTrace measurements may be attributed to two sources of variation. First, the set of cells outlined by CuticleTrace may differ from those traced by an expert in the same image. However, *CuticleTrace* generates a much larger cell measurement dataset than is possible by hand. Therefore, provided the cells selected by CuticleTrace are representative of the whole image, the differences between the two methods were more likely a result of bias in the hand-drawn cells, as the grid-selection method of hand tracing results in smaller sample sizes and tends to favor larger cells.

Second, CuticleTrace may emphasize different aspects of cell morphology than an expert. Crenulated cells with obvious three-dimensional morphology showed the greatest number of disagreements (Fig. 6a, 6d). In these images, the expert chose to outline the cell wall at an alternate focal level from the CuticleTrace macros. The choice of focal level is somewhat subjective, however, and CuticleTrace’s interpretation of the cell outline was largely reasonable and consistent. Our cell-to-cell comparison of 447 individual cells across 50 images showed that expert and CuticleTrace measurements are highly correlated (Fig. 6). For most images, CuticleTrace and expert measurements were effectively interchangeable.

The ease of data visualization built into CuticleTrace with the “Batch Overlay” and “Single Image Processor” macros is an important feature for ensuring accurate cell tracing and ROI filtering. Our design makes qualitative assessment of outputs easy, so that users can visually check all CuticleTrace outputs—thresholded images, skeletonized images, and filtered and unfiltered ROI sets—and modify input parameters as necessary. Additionally, CuticleTrace users may modify the “CuticleTrace Data Filtration” R notebook to suit their needs, and ROI sets may always be manually revised in FIJI.

Slide preparation, image quality, and CuticleTrace input parameters were the most important factors in ensuring accuracy in automated measurements. While we intentionally included a wide range of image qualities in our dataset (**Fig. S2**), we did not attempt to analyze Cuticle Database images that were of very poor quality. CuticleTrace’s flexibility allows it to be fine-tuned for a wide variety of image preparations and resolutions, but that same flexibility may lead to ineffective or inaccurate image characterization if users neglect to carefully select input parameters (Fig. 2). It is imperative that users closely follow instructions in the CuticleTrace User Manual (**Appendix S1**).

### Comparison to Alternate Automated Methods

CuticleTrace is a unique addition to the suite of available methodologies for automatically segmenting and characterizing leaf epidermal morphology. Other approaches— PaCeQuant (Möller et al., 2017) and LeafNet (Li et al., 2022)—also automate epidermal pavement cell segmentation, with some limitations. Many other methods focus on stomata detection and characterization (e.g. Fetter et al., 2019; Li et al., 2022). Additional methods have been developed to trace cells in other plant tissues (Wolny et al., 2020) and more generally (Stringer et al., 2021).

Of these methods, CuticleTrace is the first to effectively trace and measure epidermal pavement cells across a wide variety of taxa, cell morphologies, and image qualities. The customizability of the “Batch Generate ROIs” and “Single Image Processor” FIJI macros allows users to tune CuticleTrace’s settings to work well for their images. Unlike machine-learning approaches to cell segmentation, CuticleTrace does not require training images and can work with a broad range of morphologies with existing FIJI tools. It can therefore be more easily applied to studies involving diverse samples, detrital cuticle, or extinct taxa.

## CONCLUSIONS

The modularity and adaptability of the CuticleTrace toolkit hold great potential for its use outside of the immediate scope of this paper. CuticleTrace allows for the interchange of different parts of its analysis pipeline for alternate applications, and we intend for the toolkit to remain in active development. The local thresholding methods employed by CuticleTrace are effective for segmenting epidermal pavement cells in light microscopy images, but alternative segmentation methods (Möller et al., 2017; Wolny et al., 2020; Stringer et al., 2021; Li et al., 2022; Kirillov et al., 2023) can be swapped into the CuticleTrace pipeline as needed for other applications. Other methods of epidermal cell morphological analysis (Möller et al., 2017; Brown and Jordan, 2023) can also utilize CuticleTrace-generated ROIs to measure shape parameters beyond those included within CuticleTrace.

CuticleTrace’s current inability to detect and mask stomata limits its use on stomata-rich abaxial cuticle, but the toolkit’s modular structure is conducive to the integration of machine-learning-based methods for stomatal recognition and masking (Fetter et al., 2019; Aono et al., 2021; Li et al., 2022; Sai et al., 2023), which would further its applicability to abaxial leaf cuticle. With potential for integration with other methods and software, CuticleTrace represents a robust foundation for image binarization, ROI generation, and morphological analysis that may find other applications in plant science.

The leaf epidermis contains a wealth of information about plants’ evolutionary history and growth environments, and CuticleTrace makes that information significantly more accessible. In both living and fossil plants, epidermal pavement morphology can be utilized to gain insight into plant physiology, ecology, and evolution, and fossil evidence of epidermal cells are integral to some paleoenvironmental studies. The largest barrier to big-data approaches to these questions is the non-trivial task of accurately segmenting epidermal cell images. CuticleTrace represents a significant advance, greatly lowering the barrier to high-throughput studies of epidermal morphology by increasing the consistency of epidermal cell measurements and by vastly decreasing the labor investment required to generate cell shape datasets.

## ACKNOWLEDGEMENTS

RSB acknowledges financial support from NSF grant EAR:1805228. SWP acknowledges support from a 2022 University of Illinois National Center for Supercomputing Applications (NCSA) Faculty Fellowship. EDC, RED, KS, and AIM acknowledge support from NSF grant EAR:145031. BAL acknowledges financial support from Caroline Strömberg and thanks her, Alex Lowe, and Elena Stiles for constructive discussion.

## AUTHOR CONTRIBUTIONS

BAL, RSB, RED and SWP conceived the study. BAL conducted the data analysis and created the FIJI image processing macros and ‘R’ filtering code. RED, KS, and AIM provided manual cell outlines. RED and EDC recruited and supervised the University of Wyoming students. BAL and SWP wrote the manuscript with help from RSB and RED. RSB and SWP supervised and directed the research.

## DATA AVAILABILITY

All generated and analyzed data from this study are included in the published article and its Supporting Information (**Fig. S2**). The code for the FIJI macros as well as the R notebook for filtering cells is available in the GitHub repository: (https://github.com/benjlloyd/CuticleTrace).

## Supporting Information

**Appendix S1** – A detailed illustrated user manual for the CuticleTrace toolkit.

**Fig. S2** – Images of ROI outputs from CuticleTrace (Unfiltered, ±1MAD, ±2MAD) and expert hand-tracing on all 50 images in our test dataset.

