## Appendix S1 for "CuticleTrace: A toolkit for capturing cell outlines of leaf cuticle with implications for paleoecology and paleoclimatology"

The **CuticleTrace**
Image Analysis Toolkit

### User Manual

Benjamin A. Lloyd, Richard S. Barclay, Regan E. Dunn,
Ellen D. Currano, Ayuni I. Mohamaad, Kymbre Skersies,
and Surangi W. Punyasena.

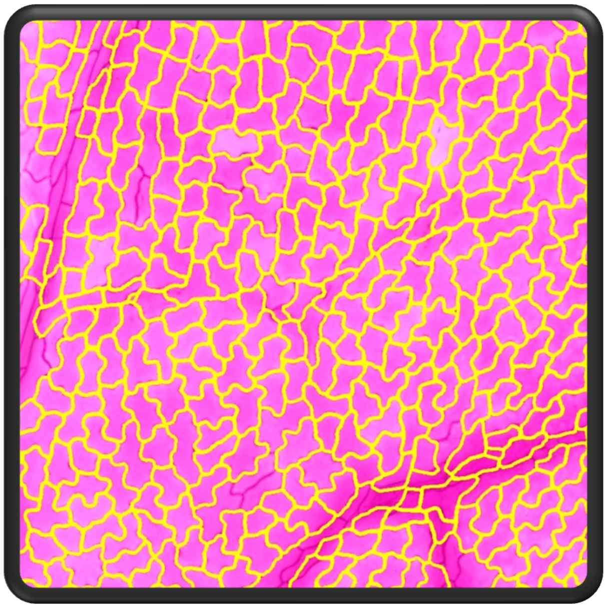

Supporting documentation for Lloyd et al., 2023:
*Cuticletrace: A toolkit for capturing cell outlines of leaf cuticle with implications for paleoecology and paleoclimatology*

### 1. Introduction

Welcome to the instructional document for the CuticleTrace image analysis toolkit. To get started, download the contents of the CuticleTrace Github repository (https://github.com/benjlloyd/CuticleTrace).

After downloading, you should have four files:
- This document - *CuticleTrace_UserManual.pdf*
- CuticleTrace_AllMacros.ijm
- CuticleTrace_DataFiltration.Rmd
- CuticleTrace_Example.zip

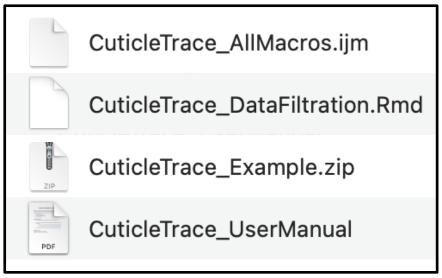

**Figure 1.** Contents of CuticleTrace GitHub repository.

The CuticleTrace toolkit consists of two files: *CuticleTrace_AllMacros.ijm* and *CuticleTrace_DataFiltration.Rmd.* To learn how to use the CuticleTrace toolkit, use the tutorial in section 2 to analyze the images contained in *CuticleTrace_Example.zip*.

#### 1.1. Toolkit contents

The CuticleTrace toolkit consists of four FIJI macros and an R notebook, all of which are described in detail in section 2.1. The contents of the CuticleTrace toolkit are as follows:

1. Four FIJI Macros. All FIJI Macros are included in *CuticleTrace_AllMacros.ijm,* consisting of:
   a. *CuticleTrace - Batch Generate ROIs*
   b. *CuticleTrace - Single Image Processor*
   c. *CuticleTrace - Batch Measure (Different Scales)*
   d. *CuticleTrace - Batch Overlay*
2. One R Notebook. Regions of interest (ROIs) generated by the CuticleTrace FIJI macros are filtered with functions in *CuticleTrace_DataFiltration.Rmd*

#### 1.2. Software requirements

CuticleTrace works on all operating systems that support FIJI and R. To install the necessary software and plugins, follow instructions below:

1. Install FIJI. FIJI is identical to ImageJ and ImageJ2, but with many plugins pre-installed. Download the software here: https://fiji.sc. This analysis pipeline was developed on FIJI running ImageJ2 version 2.9.0/1.54b.
2. Install the “Shape Smoothing” FIJI plugin from the BioMedGroup (Technical University of Dortmund, Germany) update site. Use the “Manage update sites”

menu in the FIJI updater. Follow the instructions here: https://imagej.net/update-sites/following.
3. Install R and RStudio. Download both here: https://posit.co/download/rstudio-desktop/. This analysis pipeline was developed using RStudio version 2023.03.1+446, running R version 4.2.2.

#### 1.3. FIJI macro installation

To begin analyzing images with CuticleTrace, install the FIJI Macros by following the steps below.

1. Open FIJI.
    a. Install the shape smoothing plugin if not already installed (see above).
2. Open *CuticleTrace_AllMacros.ijm* in ImageJ (drag and drop).
3. Open your Startup Macros (Process -> Macros -> Startup Macros…).
4. Select the entire contents of *CuticleTrace_AllMacros.ijm*, copy and paste at the bottom of your Startup Macros.
5. Restart FIJI. There are now 4 new macros available (Process -> Macros) (Fig. 2):
    a. *CuticleTrace - Batch Generate ROIs*
    b. *CuticleTrace - Single Image Processor*
    c. *CuticleTrace - Batch Measure (Different Scales)*
    d. *CuticleTrace - Batch Overlay*

Once FIJI has been restarted and the CuticleTrace macros appear in the Plugins menu (Fig. 2), CuticleTrace now ready for use. To learn how to use the macros, proceed to the tutorial below.

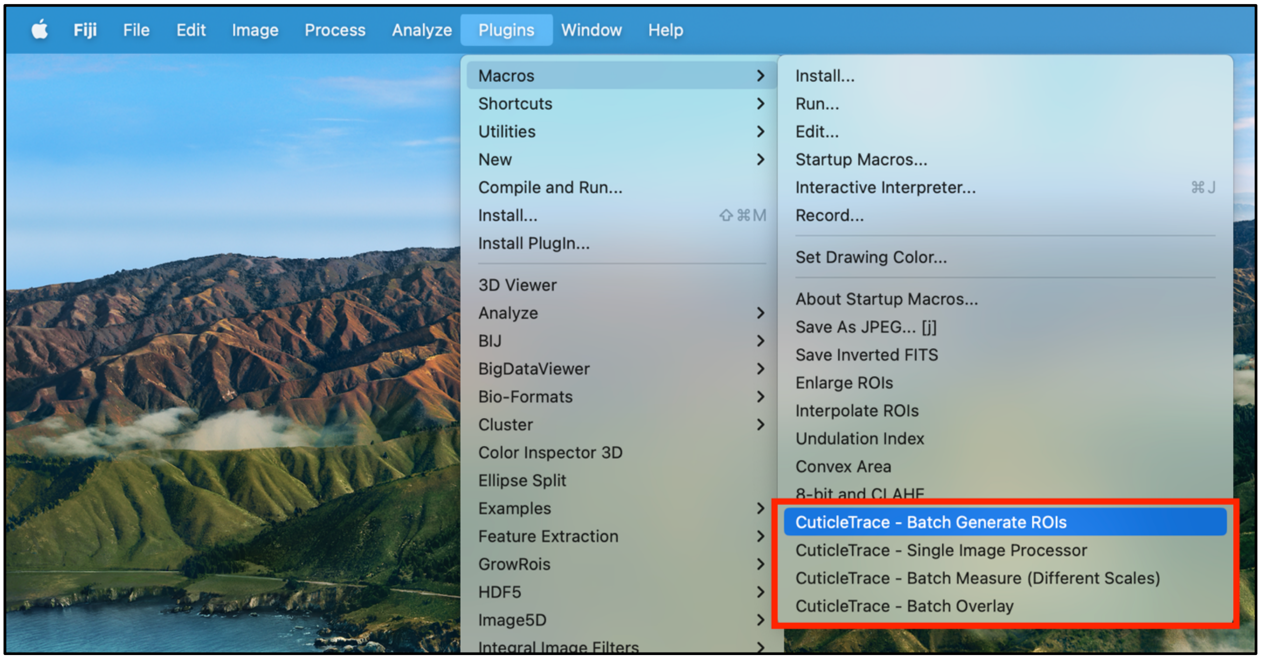

**Figure 2.** CuticleTrace FIJI macros after installation.

### 2. CuticleTrace Overview

This section provides an in-depth description of the components of the CuticleTrace toolkit (section 2.1) and running the CuticleTrace image analysis (section 2.2). To start using CuticleTrace on the example dataset (downloadable at https://github.com/benjlloyd/CuticleTrace), skip to section 3.

#### 2.1. Description of Toolkit Contents

##### Batch Generate ROIs

The "Batch Generate ROIs" FIJI macro produces thresholded and skeletonized binary images of all images in a directory, as well as the unfiltered (enhanced and interpolated) ROI sets for all images (section 3.3.1). If all images in a directory are imaged at the same scale, users may also elect to output measurements of ROI shape parameters for all images in a dataset.

##### Single Image Processor

The "Single Image Processor" FIJI macro runs the functions included in "Batch Generate ROIs" on a single image, while visualizing each step. It is used to determine input parameters for batch processing (section 3.1). It increases the transparency of CuticleTrace's image processing protocols and allows users to easily visualize the results of different macro input parameters. Furthermore, the modularity of this macro allows users to incorporate individual functions into user-generated code and scripts for specialized applications.

**Batch Measure (Different Scales)**
The “Batch Measure (Different Scales)” FIJI macro allows users to measure ROI sets generated by “Batch Generate ROIs”, if the images in a batch are of different scales (section 3.3.2). It requires that users input a CSV file that includes each image’s name and its corresponding scale. Once measurement files have been generated (either with “Batch Generate ROIs” or with “Batch Measure (Different Scales)”) unfiltered ROI sets and measurement files may be brought directly into the “CuticleTrace Data Filtration” R Notebook for statistical filtration.

**Batch Overlay**
The “Batch Overlay” FIJI macro allows users to visualize the outputs of the CuticleTrace toolkit by creating images overlain with outlines of each ROI in an ROI set, formatted to the user’s preference (e.g., Fig. 1g-i). “Batch Overlay” may be used visually check the accuracy of ROIs resulting from different input parameters and to compare the unfiltered ROI sets outputted by “Batch Generate ROIs” with the filtered versions of those ROI sets after passing them through the “CuticleTrace Data Filtration” R Notebook.

**CuticleTrace Data Filtration**
The “CuticleTrace Data Filtration” R notebook filters ROI sets with median statistics to remove erroneous ROIs. It does this by connecting the measurement files produced by FIJI to the individual file names of each ROI in each image. Then, it filters the measurements and creates new ROI sets of the remaining ROIs. It excludes measurements outside one or two median absolute deviations (MADs) from the median for area, perimeter, circularity, aspect ratio, roundness, and solidity. This approach retains ROIs that capture the central tendency of each specimen’s epidermal pavement cell morphology.

#### 2.2. Description of Processing Steps

##### 2.2.1. Image Segmentation

**Local contrast enhancement and gaussian blur**
The “Batch Generate ROIs” and “Single Image Processor” macros begin the process of image segmentation by preparing an image for thresholding. First, the image is converted to grayscale. Next, the local contrast of the image is enhanced using Contrast Limited Adaptive Histogram Equalization (CLAHE) (Zuiderveld, 1994). Then, the macros apply a Gaussian blur to the image using the specified standard deviation. This low-pass filter removes image artifacts, denoises the image, and improves local

thresholding (Misra & Wu, 2020). In setting the standard deviation of the Gaussian function, the user specifies how much to blur the image. The default value is two pixels but may need to be increased for grainy and/or noisy images (see section 3.1).

**Local thresholding**

The successful delimitation of cell walls and cell interiors forms the basis for accurate cell measurements. Thresholding transforms a grayscale image into a binary one that clearly defines individual cells (Fig. 1d). Local thresholding algorithms determine a pixel's binary value based on neighboring pixel values within a set radius. The selection of thresholding algorithm and initial radius is extremely important for generating accurate cell outlines. The choice of algorithm and initial radius will vary with imaging method and image resolution, so experimentation is necessary before batch-processing (see section 3.1). The "Batch Generate ROIs" and "Single Image Processor" macros use FIJI's native "Auto Local Threshold" plugin (Schindelin *et al.*, 2012) and allow users to select any of FIJI's built-in local thresholding algorithms (Otsu, 1979; Niblack, 1986; Sauvola & Pietikäinen, 2000; Soille, 2004; Sankur, 2004; Phansalkar *et al.*, 2011; Schindelin *et al.*, 2012; Vatani & Enayatifar, 2015).

The initial threshold radius is set by the user and iteratively refined by CuticleTrace. Each new iteration's thresholding radius is equal to the average minor radius of the fit ellipses of all cell interiors. Calibrating the thresholding radius to the cell size preserves cell shape. Too large of a radius results in artificially thickened and inaccurate cell walls, while too small of a radius ineffectively segments the image. In an image set with relatively uniform cell dimensions, a single thresholding radius may be selected and iterative thresholding may not be necessary. Choosing a single thresholding radius reduces image processing time.

**Shape smoothing and skeletonization**

Thresholding results in a binary image, with cell interiors as white objects on a black background. The "Batch Generate ROIs" and "Single Image Processor" macros then smooth the resulting cell interior shapes using the "Shape Smoothing" FIJI plugin (Wagner, 2016). The plugin uses discrete Fourier transformation to produce Fourier descriptors of all image contours. The Fourier descriptors are then filtered by relative percentage to produce smoothed contours (Wagner, 2016) (Fig. 1e). The CuticleTrace default is to retain 5% of Fourier descriptors, but the value should be empirically determined for each input dataset.

Next, the "Batch Generate ROIs" and "Single Image Processor" macros reduce cell walls to a uniform thickness using FIJI's native skeletonization algorithm (Zhang & Suen, 1984). Skeletonization distills the thresholded image to a single-pixel network that captures the outlines of epidermal cell shape. The macros then widen the skeletonized outline to three pixels for cell measurements (Fig. 1f).

##### 2.2.2 Image Analysis

**Particle analysis - ROI detection and correction**

Once we have a binary, skeletonized image representing the cell outlines, the “Batch Generate ROIs” and “Single Image Processor” macros utilize FIJI’s Particle Analysis tool to identify all discrete objects in a binary image and outline them as ROIs (Schindelin *et al.*, 2012). In our case, the ROIs are whole epidermal cells (the desired output) and partial or connected cells where the cell walls were not accurately characterized during segmentation (an unwanted output).

Inaccuracies in thresholding and skeletonization are the result of variations in brightness and contrast within an image. This sometimes results in skeletonization artifacts such as short line segments that intrude into cell interiors (Fig. 1g). The “Batch Generate ROIs” and “Get ROIs from Skeletonized Image” macros remove these errors by enlarging ROIs to fill incursions, then reducing the enlarged ROIs so that the cell outline falls in the center of the cell wall, with a one-pixel gap between cells (Fig. 1h). The resulting ROI sets minimize inaccurate cell tracings, but still may include partial-cell or multi-cell ROIs that must be removed from the final dataset (Fig. 1h). We filter out these ROIs and measurements at the end of our workflow, before visualizing results (***ROI filtration*)**.

The “Batch Generate ROIs” and “Single Image Processor” macros also use FIJI’s interpolation function to smooth the ROI outline and better capture the true cell shape. Interpolation uses a three-point running average, then converts each ROI into a polygon with vertices spaced at a defined interval (Schindelin *et al.*, 2012). The macros calculate the interpolation interval as 1/100 of the circumference of a circle with an area equal to the average image ROI area. This allows the interpolation intervals to be objectively scaled with ROI pixel dimensions across multiple images.

**ROI measurement**

The “Batch Generate ROIs” macro can also generate measurements of all ROIs of each image in a batch, provided that all images are at the same scale. The “Batch Measure (Different Scales)” macro allows users to generate measurements of all ROIs of each image in a batch if images are at different scales. The “Single Image Processor” macro may be used to measure ROIs from a single image. All three macros utilize FIJI’s native “measure” function to generate these results and write the results to CSV files that are subsequently saved into the specified output directory. The results files contain 11 shape measurements for each ROI (Table 1).

**Table 1.** Descriptions or equations of all shape parameters measured by CuticleTrace.

| Measurement Parameter | Description or Equation |
|---|---|
| Area | The area of an ROI. |
| Perimeter | The length of the outside boundary of an ROI. |
| Bounding Rectangle | The X-Y coordinates and dimensions of the smallest rectangle that encloses an ROI. |
| Fit Ellipse | The dimensions and orientation of an ellipse fit to an ROI. |
| Feret's Diameter | The longest distance between any two points on the ROI boundary. |
| Minimum Feret's Diameter | The shortest distance between any two points on the ROI boundary. |
| Circularity | $4\pi * \frac{[Area]}{[Perimeter]^2}$ |
| Aspect Ratio | $\frac{[Major\ Axis\ of\ Fit\ Ellipse]}{[Minor\ Axis]}$ |
| Roundness | $4 * \frac{[Area]}{\pi * [Major\ Axis\ of\ Fit\ Ellipse]^2}$ |
| Solidity | $\frac{[Area]}{[Area\ of\ Convex\ Hull]}$ |
| Undulation Index | $\frac{[Perimeter]}{2\pi * \sqrt{[Area]/\pi}}$ |

#### ROI filtration

The ROIs produced by the CuticleTrace FIJI analysis pipeline include partial cells, multiple cells, vein cells, or non-cuticle image artifacts (e.g., slide background, debris). The final step is to remove these erroneous ROIs. We use RStudio (Posit Team, 2023) to take advantage of R's (R Core Team, 2022) data formatting capabilities. The "CuticleTrace Data Filtration" R notebook removes ROIs with measurements outside one or two median absolute deviations (MADs) from each image's median value for area, perimeter, circularity, aspect ratio, roundness, or solidity (Table 1). We use the median as our reference point, as unfiltered measurements are non-normally distributed. "CuticleTrace Data Filtration" produces new, filtered results files and ROI sets that can be opened directly in FIJI and overlaid over the original image (Fig. 1i).

**Visual check of filtered data**

Visualizing CuticleTrace outlines is an important step in verifying that the ROIs retained after filtering accurately characterize a representative subset of epidermal pavement cells. The “Batch Overlay” macro simplifies the process of visualizing ROI sets across many images by generating new image sets with overlaid ROIs that are formatted for optimal viewing.

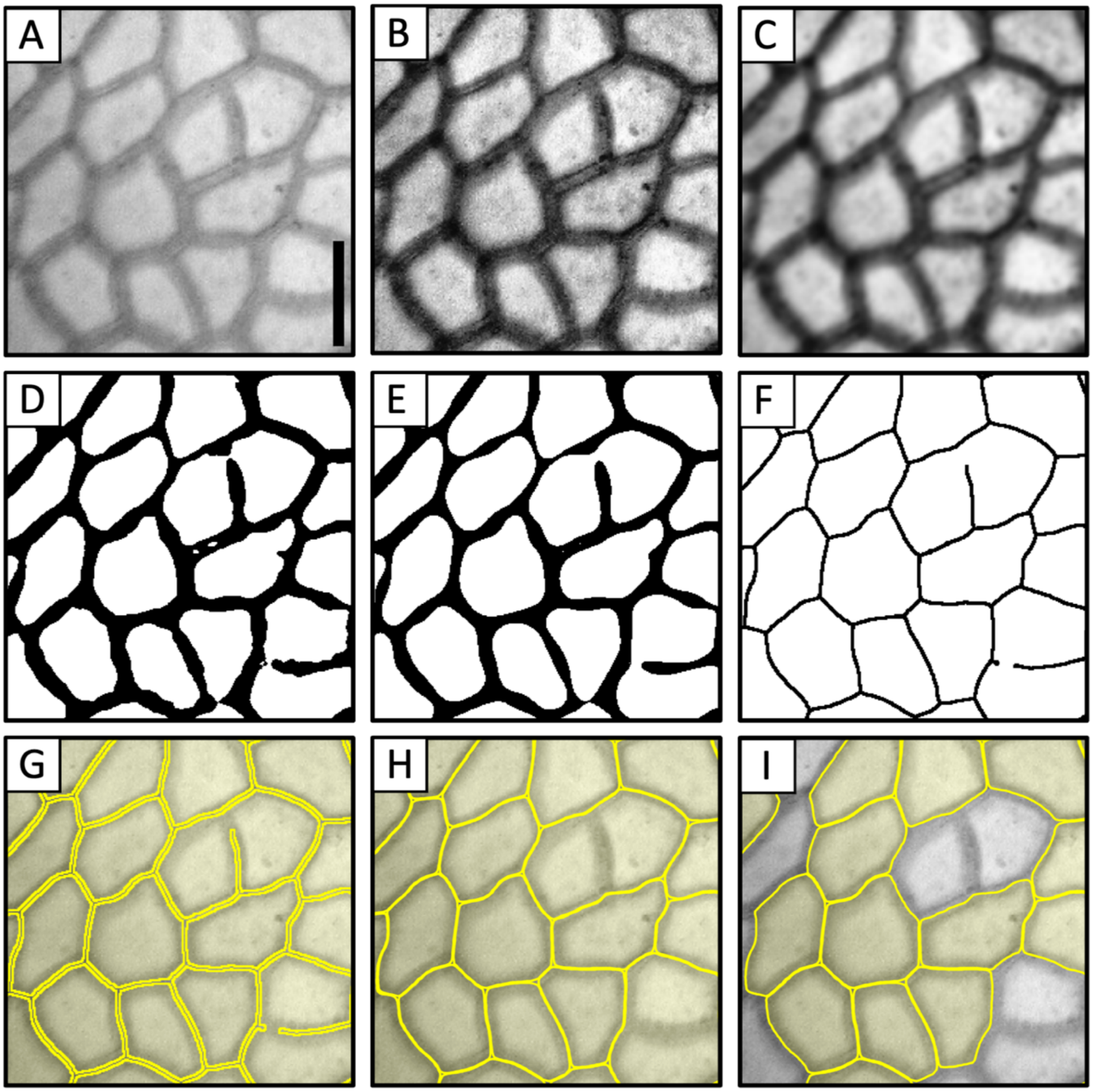

**Figure 3.** Processing steps for the CuticleTrace toolkit, illustrated on an image of **(A)** *Castanea pumila* (Fagaceae, FLMNH00115). Scale bar is 10 µm. Processing starts with segmenting epidermal cells by **(B)** enhancing local contrast, **(C)** applying gaussian blur, **(D)** local thresholding, **(E)** shape smoothing, and **(F)** skeletonization. Particles are analyzed in FIJI, by **(G)** generating ROIs, **(H)** enlarging and interpolating ROIs, and measuring ROIs. ROI measurements are **(I)** filtered with median statistics in R.

### 3. Tutorial - Analysis of example dataset

In this tutorial, we will generate final ROI sets and epidermal pavement cell measurements of the 10 images included in the CuticleTrace_Example.zip file.

#### 3.1. Example dataset contents

The example dataset provided in the CuticleTrace GitHub repository is stored as a .zip file. Once expanded, it contains two folders:

1. *Example_Dataset* - This folder contains 10 cropped images from the Cuticle Database, and a CSV file with each image's scale. We will process them in the coming steps (Fig. 4).
2. *Example_Dataset_COMPLETE_ForComparison* - This is a folder of all the results that can be generated with the images in the *Example_Dataset* folder. It is there for comparison, to make sure that you know how things are supposed to look.

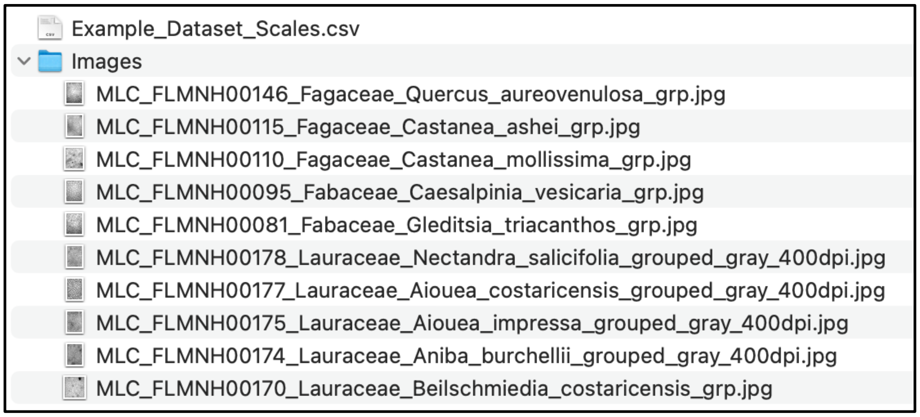

**Figure 4.** Contents of *Example_Dataset*, showing a folder of 10 images and a CSV file with each image's scale.

#### 3.2. FIJI macro input parameters

The first step in the generation of an epidermal pavement cell shape dataset with CuticleTrace is the determination of input parameters for batch image processing. **This is a non-trivial process and is necessary for the generation of accurate data.** For this tutorial, we have predetermined ideal input parameters for the example dataset (Table 1, Fig. 5). To determine the ideal input parameters for other datasets using "Single Image Processor", see section 4.1 (Determination of Input Parameters).

**Table 1.** Ideal batch processing inputs for the example image set, as determined by Lloyd et al. (2023).

| Input Parameter | Value |
|---|---|
| Cell Walls on Background | Dark on Light |
| Gaussian Blur | 2 pixels |
| Thresholding Method | Sauvola |
| Initial Thresholding Radius | 50 pixels |
| ROI Size Filter | 500-50,000 pixels$^2$ |
| Smoothing Value | 5% FDs retained |
| Scale | Various (see 3.3.2) |

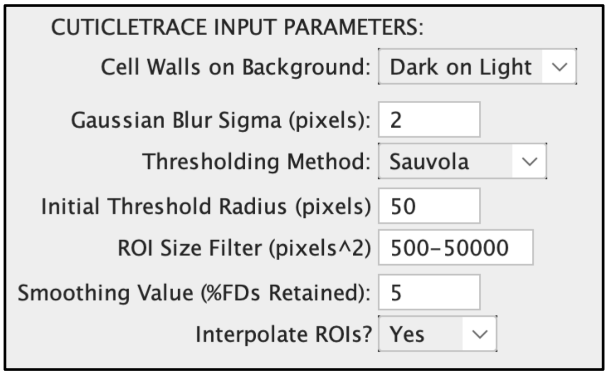

**Figure 5.** CuticleTrace Input Parameters with correct values for the example dataset, as they appear in the "Batch Generate ROIs" FIJI macro.

#### 3.3. Batch processing in FIJI

The second step in the generation of an epidermal pavement cell shape dataset with CuticleTrace is the batch processing of images with the "Batch Generate ROIs" and "Batch Measure (Different Scales)" macros in FIJI. This step generates unfiltered ROI sets and results files that include accurate cell tracings as well as some undesired ROIs. Ideal batch processing inputs for the example dataset are in Table 1 and Fig. 5.

##### 3.3.1. Batch Generate ROIs

With your input parameters selected and your output directories created, you are ready to batch process the example image set with the "Batch Generate ROIs" macro.

1. In FIJI, with no images open, open the "Batch Generate ROIs" macro (Plugins > Macros > CuticleTrace - Batch Generate ROIs)
2. First, a "NOTICE" window will pop up, reiterating important points regarding the use of the macro. To continue, click OK.
3. The next window contains the input fields necessary to run the macro:
    a. Select the folder containing the image set for the input directory (Fig. 4).
    b. Double-check that the parameters in Table 1 match the default inputs in "Batch Generate ROIs" (Fig. 6).
    c. As is evident in *Example_Dataset_Scales.csv*, not all these images have the same scale. **Therefore, do not select "generate results files"**.
    d. Click OK.

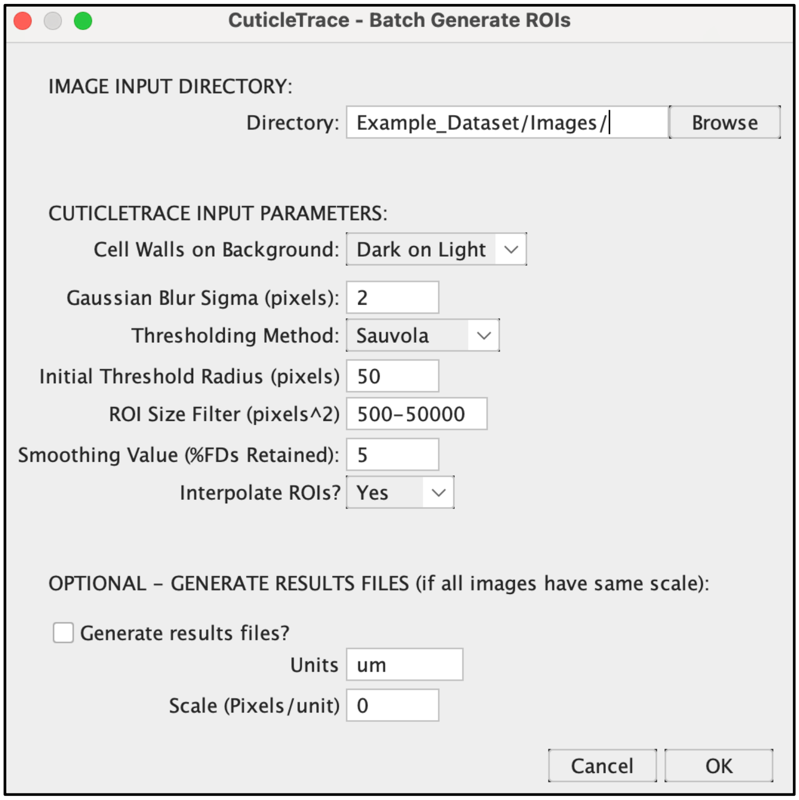

**Figure 6.** The input window of "Batch Generate ROIs" with the correct inputs to batch process the example dataset.

"Batch Generate ROIs" will run for a few minutes—you should see images popping up sequentially. Once it is finished, you will have new folders adjacent to your Image Folder, each containing the macro's outputs:

- *Thresholded_Images* should contain thresholded and smoothed binary images of each of the 10 input images (Fig. 7c).
- *Skeletonized_Images* should contain skeletonized versions of the thresholded images (Fig. 7d).
- *ROI_Sets* should include 10 zipped ROI Sets, 1 for each image (Fig. 7e).
- *Image_Metadata* should include 10 small CSV files with the thresholding radius and interpolation interval of each image (Fig. 7b), as well as 1 larger CSV file containing all the thresholding radii and interpolation intervals for the dataset**. We recommend adding these data into *Example_Dataset_Scales.csv*, to keep scales, thresholding radii, and interpolation intervals together in one dataset.**

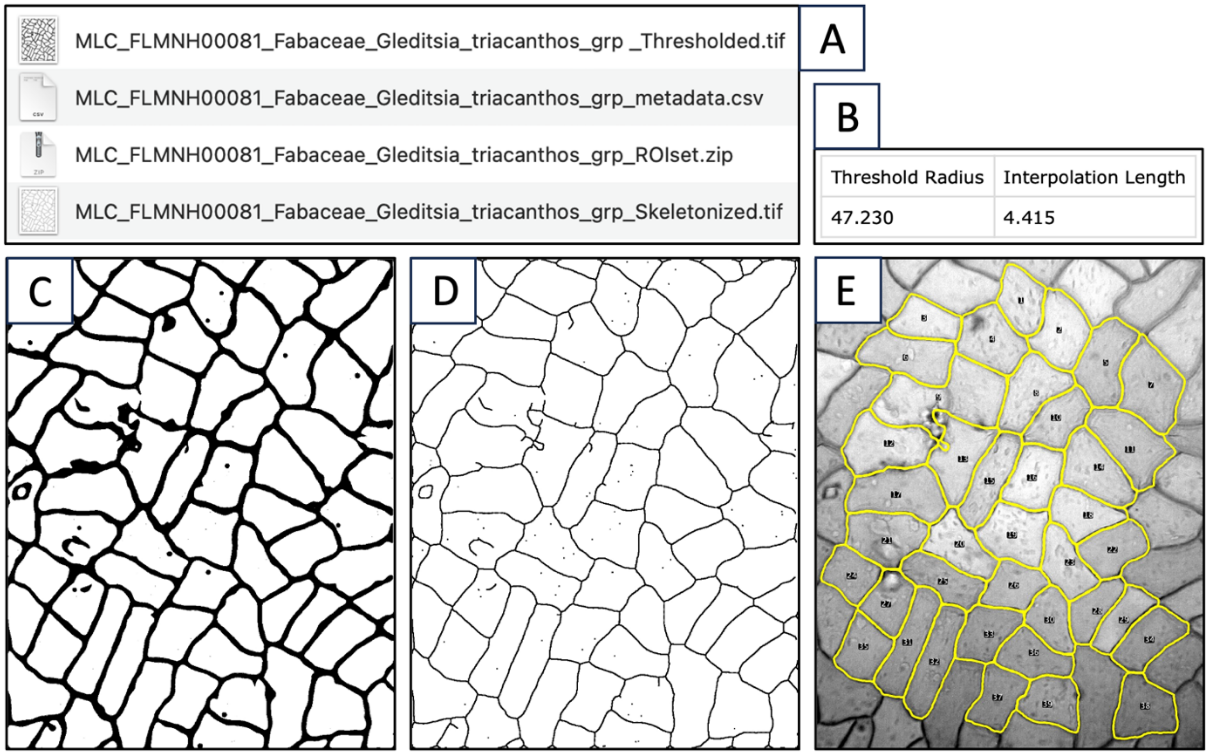

**Figure 7.** The outputs of “Batch Generate ROIs” from one image (FLMNH00081, Gleditsia triacanthos). **(A)** The four files output by the macro, including **(B)** the image metadata, **(C)** the thresholded image, **(D)** the skeletonized image, and **(E)** the unfiltered ROI set (shown overlaying the original image).

##### 3.3.2. Batch Measure (Different Scales)

Once you have verified that “CuticleTrace - Batch Generate ROIs” worked correctly, it is time to measure your ROI sets. Since not all these images are the same scale, we use “CuticleTrace - Batch Measure (Different Scales)”*.* This macro requires a CSV file with a column for the image names, and a column with the scale of each image. For the example dataset, this is *Example_Dataset_Scales.csv*.

1. In FIJI, with no images open, open the “Batch Measure (Different Scales)” macro (Plugins > Macros > CuticleTrace - Batch Measure (Different Scales))
2. Select *Example_Dataset_Scales.csv* as the CSV file, and double check that the column names match the default text (they should). Then select the folders containing your unprocessed images and your ROI sets folder.
3. Click OK.

“Batch Measure (Different Scales)” will run for a few minutes but should be much faster than “Batch Generate ROIs.” You should see images popping up in quick succession. Once it is finished, you will have a new folder, *Results_Files*, containing 10 new CSV files (Fig. 8). When you open these files, you should see measurements for all shape parameters discussed in Lloyd et al. (2023).

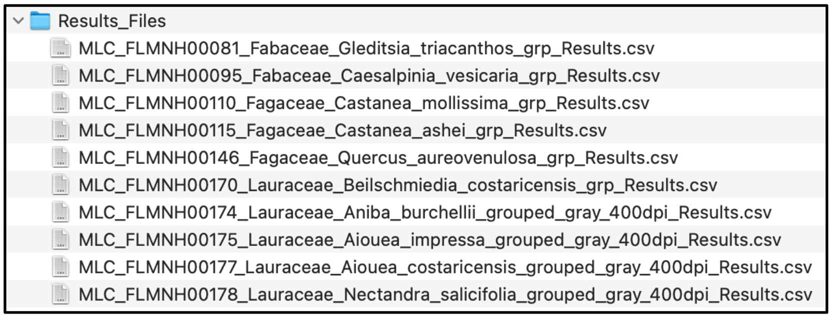

**Figure 8.** Results files as they appear after running “Batch Measure (Different Scales)”.

#### 3.4. Data filtration in R

Now that you have generated your unfiltered datasets, it is time to remove inaccurate and unwanted ROIs by filtering ROI sets with median statistics. *CuticleTrace_DataFiltration.Rmd* creates new ROI sets and results files that have been filtered to ±1 MAD and ±2 MAD of the median of area, perimeter, circularity, Feret’s diameter, aspect ratio, roundness, and solidity. For more detail on ROI filtering specifics, see the main text of Lloyd et al. (2023).

##### 3.4.1. CuticleTrace_DataFiltration.Rmd

To begin the filtering process, copy CuticleTrace_DataFiltration.Rmd into the Example_Dataset folder. **To work correctly, CuticleTrace_DataFiltration.Rmd must be in the same directory as your *Results_Files* and *ROI_Sets* folders.** Follow these steps to filter your data and ROI sets:

1. Open CuticleTrace_DataFiltration.Rmd in RStudio. You will see sections of text detailing the filtering procedure broken up by code chunks. Code chunks may be collapsed for increased readability.
2. Read through the filtering procedure.
3. Make sure that the file paths that lead to your *Results_Files* folder (line 361) and *ROI_Sets* folder (line 367) are correct (Fig. 9). If your folders are named “Results_Files” and “ROI_Sets” and are in the same folder as *CuticleTrace_DataFiltration.Rmd* (as recommended), the default text will work.
4. The code can be run in two ways:
    a. As you read through the filtering procedure, you may run each code chunk as you get to it by clicking the green triangle on the upper right corner of each code chunk (Fig. 9). This is a more transparent way to run the code and is recommended for a first-time user.
    b. If you want to move quickly, click Knit (Knit to HTML) on the toolbar. This will run the entire code and create a nicely formatted HTML version of the notebook to be saved alongside the .Rmd file.

```
356 1.  Make sure this .Rmd file is within the directory that contains your
    Results folder and your ROI sets folder. .Rmd files automatically set the
    working directory to the directory that contains the document. **If using
    the example dataset, place this .Rmd file within the "Example_Dataset"
    folder.**
357
358 2.  Paste the directory path that contains your results files.
359
360 ```{r}
361 results_path <- "Results_Files/"
362 ```
363
364 3.  Paste the directory path that contains your ROI sets.
365
366 ```{r}
367 ROIsets_path <- "ROI_Sets/"
368 ```
```

**Figure 9.** Line 361 and line 367 of CuticleTrace_DataFiltration-.Rmd must lead to the Results and ROI Sets directories.

**Now, you have generated your final, filtered datasets.** Navigate to your *Results_Files* (Fig. 10) and *ROI_Sets* (Fig. 11) folders to see the results.

- In your *Results* folder, you should see a new *Results_Reformatted* folder, which will contain a reformatted version of the unfiltered datasets. It will also contain two additional folders:
    - *1MAD_Trimmed_Results* contains results files trimmed to within 1 MAD of the median of area, perimeter, circularity, Feret's diameter, aspect ratio, roundness, and solidity.
    - *2MAD_Trimmed_Results* contains results files trimmed to within 2 MAD of the median of area, perimeter, circularity, Feret's diameter, aspect ratio, roundness, and solidity.

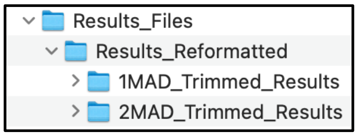

**Figure 10.** New Results_Reformatted folder within Results_Files, with directories containing ±1 MAD and ±2 MAD filtered datasets.

- In your ROI Sets folder, you should see an unzipped version of all ROI sets, as well as a zipped* and unzipped version of new ROI sets that have been filtered to ±1 MAD and ±2 MAD of the median of area, perimeter, circularity, Feret's diameter, aspect ratio, roundness, and solidity. These ROI sets correspond to each newly generated results file.

  ***NOTE: The current version of CuticleTrace_DataFiltration.Rmd does not generated zipped ROI sets on Windows machines. To view filtered ROI sets in FIJI, manually zip them prior to opening them in FIJI (section 3.5).**

##### 3.4.2. Organization of filtered ROI sets

We recommend reorganizing the ROI set outputs of *CuticleTrace_DataFiltration.Rmd* into 3 new folders within the *ROI Sets* folder, each with a *Zipped* and *Unzipped* folder within (Fig. 11). Refer to the folder *Example_Dataset_COMPLETE_ ForComparison* for an example.

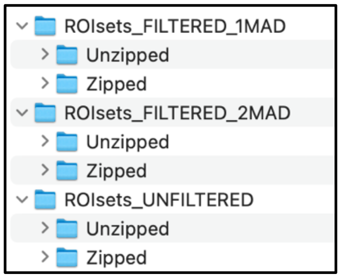

**Figure 11.** Recommended file organization for filtered ROI sets.

#### 3.5. Visualize results in FIJI

To visually inspect the results of CuticleTrace ROI generation and filtering, it is important to create new image sets showing filtered and unfiltered ROI sets for each image. To do this, you may utilize the “CuticleTrace – Batch Overlay” macro in FIJI. Resulting images should look generally like the image in Fig. 7e.

##### 3.5.1. Batch Overlay

Open FIJI and select the “Batch Overlay” macro (Plugins > Macros > CuticleTrace - Batch Overlay). Select the original, unprocessed image folder (*Images*) as the image input directory, and the ***Zipped*** Unfiltered ROI set folder as the ROI set directory. **FIJI cannot read unzipped ROI sets.** Since we are starting with the Unfiltered dataset, name the output folder “Images_with_Overlays_ Unfiltered” or similar (Fig. 12).

“Batch Overlay” contains a lot of options for changing line thickness, fill color, and more, but we recommend leaving the inputs at their default values for the example dataset. Once you have selected the image directory, ROI set directory, and output folder name, click OK.

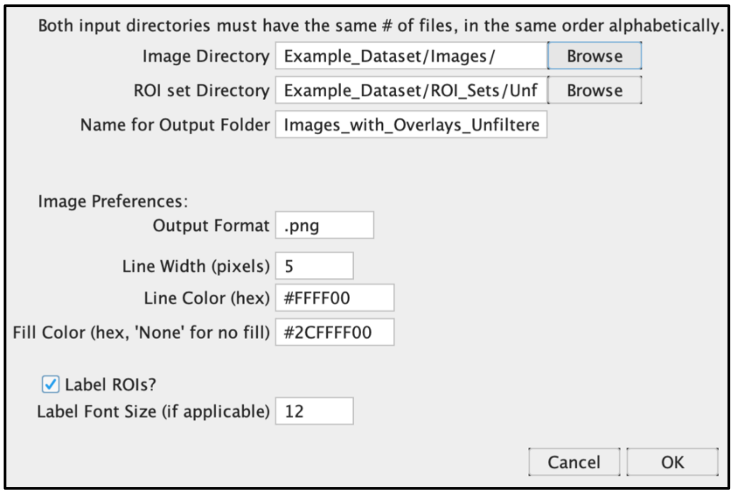

**Figure 12.** Correct inputs for processing Example Dataset images with the “Batch Overlay” FIJI macro.

“Batch Overlay” will run for a few minutes—you should see images flashing across the screen. Once it is complete, run it again with the ±1 MAD and ±2 MAD ROI set input and output directories. Once this is complete, you may navigate to these folders and view your final CuticleTrace-generated ROI sets overlain on their source images. Good work! Since you now have 3 folders of Images with Overlays, we recommend reorganizing them into their own directory, containing 3 folders for images overlain with unfiltered, ±1 MAD, and ±2 MAD ROI sets (Fig. 13).

You may now compare these images to those in the *Example_Dataset_COMPLETE_ForComparison* folder to verify that you generated the same results.

Congratulations! You have now used the full capability of the CuticleTrace toolkit.

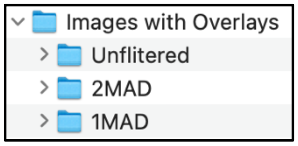

**Figure 13.** Recommended folders for “Batch Overlay” outputs.

### 4. Using CuticleTrace with new datasets

Once you have successfully generated and measured ROIs from our provided example dataset, it’s time to move on to your own images. The level of customization available within CuticleTrace’s input parameters allows this toolkit to be applied to a wide variety of cell morphologies and image qualities. **However, the flexibility built into CuticleTrace means it is especially important to determine appropriate input parameters for new image sets.** Below, we detail how to use FIJI’s native functionality and the “CuticleTrace – Single Image Processor” macro to determine ideal input parameters for your dataset.

#### 4.1. Determination of input parameters

The first step in the generation of an epidermal pavement cell shape dataset with CuticleTrace is the determination of input parameters for batch image processing.

##### 4.1.1 Overview of input parameters

The input parameters required to batch process images are covered in depth in section 2.2. Input parameters may be seen as they appear in the CuticleTrace FIJI user interface in Table 3.

For images to be batch processed, all input parameters (except scale) **must be identical**. Images of different scales may be analyzed in the same batch by utilizing the

“Batch Generate ROIs” macro to generate ROIs, followed by measurement with the “Batch Measure (Different Scales)” macro.

**To efficiently and effectively batch process image sets, it is necessary to experiment with a subset of images to determine the ideal input parameters for the “Batch Generate ROIs” macro.**

**Table 3.** Input parameters for CuticleTrace FIJI Macros.

| Input Parameter | Options/Default Value | Method of Determination |
|---|---|---|
| Cell Walls on Background | *Dark on Light, Light on Dark* | Determined visually |
| Gaussian Blur (pixels) | *2 pixels* | Determined experimentally on a subset of images with “CuticleTrace - Single Image Processor” (section 3.1.3) |
| Thresholding Method | *Bernsen, Contrast, Mean, Median, MidGrey, Niblack, Otsu, Phansalker, Sauvola* | Determined experimentally on a subset of images with “CuticleTrace - Single Image Processor” and *Auto-Local-Threshold* (section 3.1.3) |
| Initial Threshold Radius (pixels) | *50 pixels* | Determined by measuring a subset of approximate cell widths (section 3.1.2) |
| ROI Size Filter (pixels$^2$) | *500-50,000 pixels$^2$* | Determined by measuring a subset of approximate cell areas (section 3.1.3) |
| Smoothing Value (%FDs retained) | *5% FDs retained* | Determined experimentally on a subset of images with “CuticleTrace - Single Image Processor” (section 3.1.3) |
| Scale (pixels/unit) | No default value * | Should be known before processing |

* One single value may be entered in the “Batch Generate ROIs” macro if all images are the same scale, or a CSV file specifying the scale of each image may be imported with “Batch Measure (Different Scales)”.

#### 4.1.2. Initial Thresholding Radius

The initial thresholding radius to apply to an image set is best determined by approximating a subset of cell widths. **The ideal initial thresholding radius is approximately half of the average pavement cell width in the whole dataset.** It is iterated upon for every image, so it does not have to be exact. To determine an ideal initial thresholding radius, we recommend measuring the approximate widths of ~10 cells across ~5 images (1-3 cells per image), to get a reasonable average value to apply to the dataset (Fig. 14).

- To measure approximate cell width (in pixels), remove the image scale (Analyze > Set Scale… > "Click to Remove Scale"). Then select the straight-line tool from the toolbar and draw a line across the width of a cell (Fig. 14). Measure the line by pressing the M key. Measure ~1-2 more cells, and then repeat this process with ~3-5 other images. **Set the initial thresholding radius to approximately ½ of the average of your length measurements.**

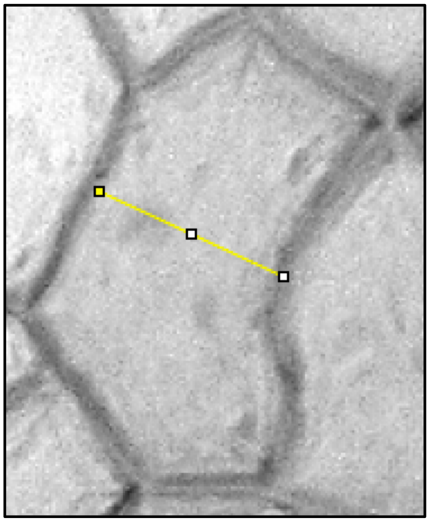

**Figure 14.** A cell from FLMNH00081 (*Gleditsia triacanthos*), measured by drawing line across its width. This cell measured 95 pixels wide.

##### 4.1.3. ROI Size Filter

The ROI size filter to apply to an image set is best determined by approximating the cell areas of the smallest and largest cells in the dataset (Fig. 15). **The ideal ROI size filter captures all cells in an image set, while removing ROIs that are much too small or large to consist of an accurately traced cell.** ROIs will be filtered further with the R notebook *CuticleTrace_DataFiltration.Rmd*, so some unwanted ROIs are expected to remain in the dataset at this step.

- To measure approximate cell area (in pixels), remove the image scale (Analyze > Set Scale… > "Click to Remove Scale"). Then select the elliptical selection tool from the toolbar and draw an oval around a cell, roughly approximating its area (Fig. 15). Measure the ellipse by pressing the M key. Measure ~2-5 more of the smallest and largest cells you observe, and then repeat this process with ~3-5 other images. **Set the ROI size filter to widely encompass the smallest and largest cells in your dataset, within an order of magnitude.**

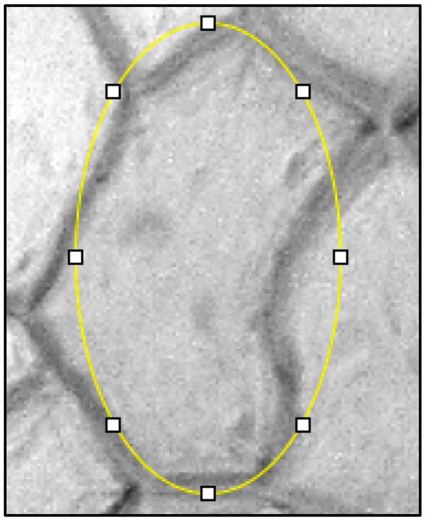

**Figure 15.** A cell from FLMNH00081 (*Gleditsia triacanthos*), measured by an ellipse of similar dimensions. This ellipse had an area of 20,080 pixels$^2$.

##### 4.1.4. Thresholding Method

The easiest way to determine which local thresholding algorithm is likely to work best for your dataset is with FIJI's built-in *Auto Local Threshold* function. To use this function, first make your image 8-bit (Image > Type > 8-bit), enhance its local contrast (Process > Enhance Local Contrast (CLAHE) > OK), and apply a default Gaussian Blur of 2 pixels (Process > Filters > Gaussian Blur…). Then, proceed to the Auto Local Threshold menu (Image > Adjust > Auto Local Threshold). Input your previously measured thresholding radius and ROI size filter and select "Try All" for the thresholding method (Fig. 16).

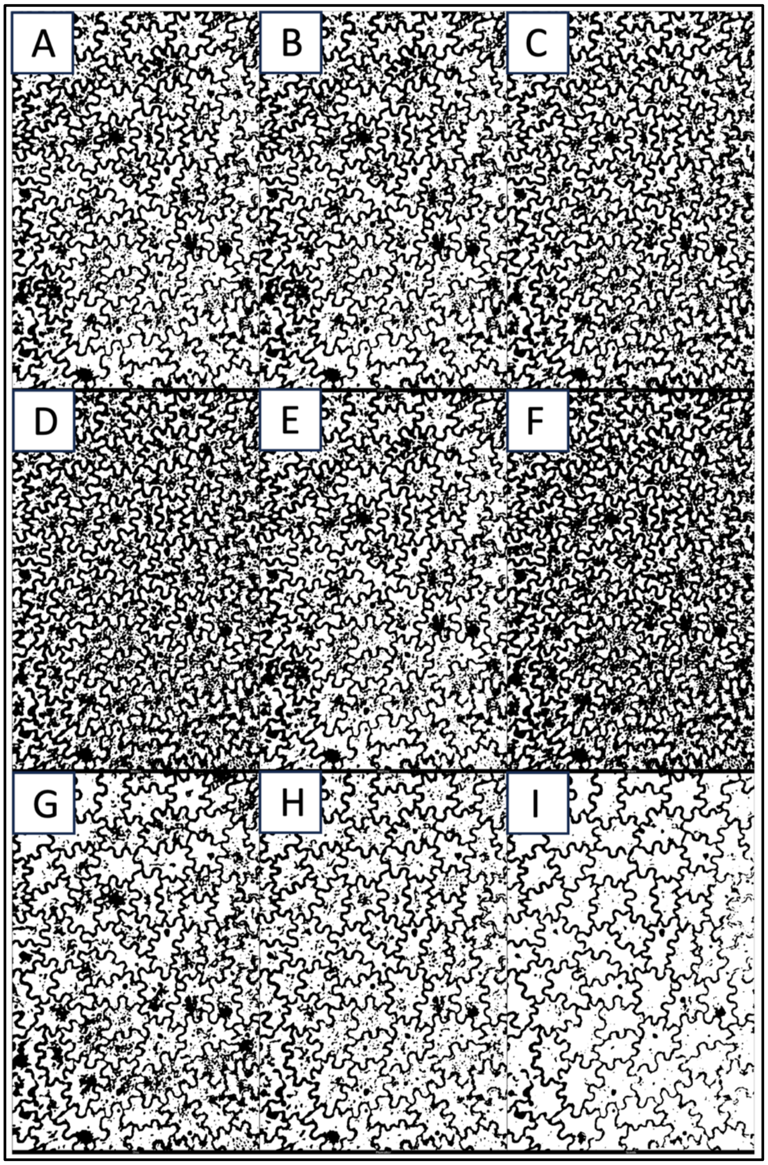

**Figure 16.** The output of FIJI's Auto Local Threshold "Try All" option on FLMNH00510 (*Toxicodendron striatum*). Thresholding methods are **(A)** Bernsen, **(B)** Contrast, **(C)** Mean, **(D)** Median, **(E)** MidGrey, **(F)** Niblack, **(G)** Otsu, **(H)** Phansalker, **(I)** Sauvola (optimal for this image).

Once you have an image montage with all local thresholding algorithms, note which methods work best for your image. These will be fine-tuned by “Batch Generate ROIs” and “Single Image Processor”, so they do not have to be perfect. For the image in Fig. 16, Sauvola thresholding (Fig. 16i) works best, and Phansalker thresholding (Fig. 16h) does acceptably. Repeat this process with ~3-5 images in your dataset to see which thresholding algorithms tend to work well for your images.

##### 4.1.5. Thresholding Method, Gaussian Blur, and Smoothing Value with “Single Image Processor”

Once you have determined your initial thresholding radius and ROI size filter, and you have narrowed down your options for thresholding method, it is time to utilize the “CuticleTrace – Single Image Processor” macro to determine the ideal thresholding method, gaussian blur value, and smoothing value for your dataset. **It is necessary to experiment on and qualitatively evaluate a subset of images to determine these values.**

“Single Image Processor” brings an open image in FIJI through all (or some of) the steps in the CuticleTrace analysis pipeline (see Fig. 1 of Lloyd et al., 2023). **It allows you to observe the combined effects of these inputs and to determine ideal inputs for your dataset.** It may also be used to individually analyze images instead of batch processing.

To use “Single Image Processor”, open the macro (Plugins > Macros > CuticleTrace – Single Image Processor). The dialog box that appears will contain options to modify all CuticleTrace input parameters, and an option to select “desired outputs” (Fig. 17). The “desired outputs” menu allows you to decide what steps from the CuticleTrace pipeline you want to visualize (Fig. 18). The default option, “Thresholded and Skeletonized Image with ROIs and Results”, brings an unprocessed image through the entire pipeline (Fig. 19). Other options include fewer steps and/or allow for the processing of a previously thresholded or skeletonized image. In Fig. 19, we show the outputs of the “Single Image Processor” at two different smoothing values, to illustrate the macro’s ability to visualize outputs.

**Using “Single Image Processor”, empirically determine the gaussian blur value, thresholding method, and smoothing value that most accurately characterize cells across a subset of images. With these values** (and your previously determined initial thresholding radius and ROI size filter) **you are ready to batch process your image set.**

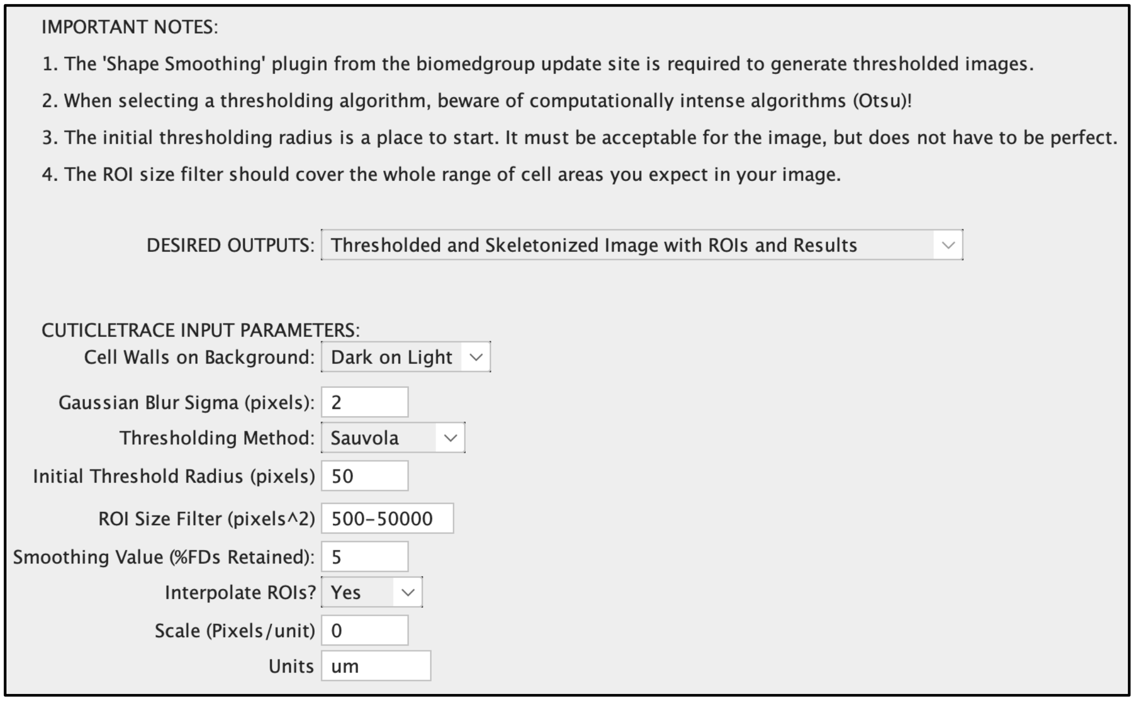

**Figure 17.** The input window for the “Single Image Processor” with default inputs for the example dataset.

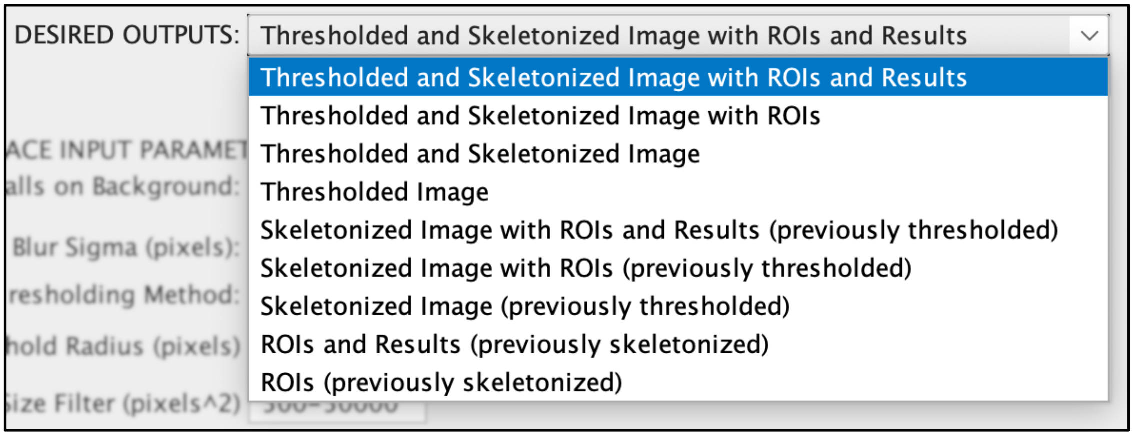

**Figure 18.** The "Desired Outputs" menu options within the “Single Image Processor”.

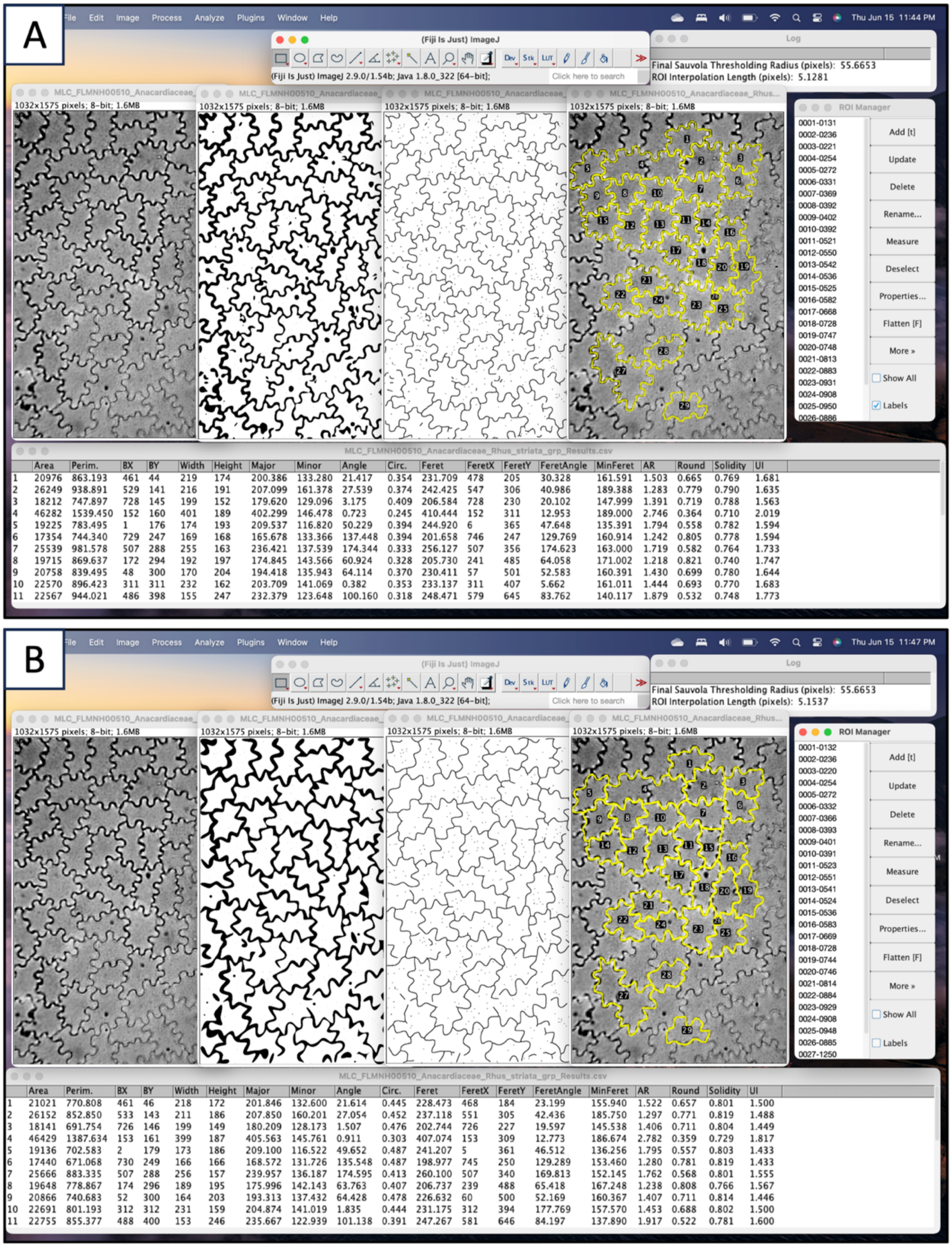

**Figure 19.** Outputs of “Single Image Processor” on FLMNH00510 (Toxicodendron striatum) with **(A)** all default input parameters (as seen in Fig. 16) and **(B)** all default input parameters **except smoothing value** (2% FDs retained). The default smoothing value in **(A)** (5% FDs retained) performs better for this image.

##### 4.1.6. Troubleshooting - Manually replicate “Single Image Processor”

If you are experiencing issues with the CuticleTrace FIJI macros, or if you would like to further investigate input parameters at a more granular level of detail than what is supported in “Single Image Processor,” follow these instructions to approximately replicate the steps of the CuticleTrace image analysis pipeline through FIJI’s user interface. **This is not necessary to determine input parameters but is provided for additional transparency.**

1. Make the image 8-bit (Image > Type > 8-bit)
2. Enhance local contrast (Process > Enhance Local Contrast (CLAHE) > OK)
3. Apply Gaussian Blur (Process > Filters > Gaussian Blur…)
    a. In this step, you may experiment with different values to see how they influence downstream results. In general, the default value of 2 pixels works well with images of similar resolution of our example dataset. In general, increased blurring helps smooth out differences in contrast along cell walls but will erode details in cell shape at higher values (Fig. 20).

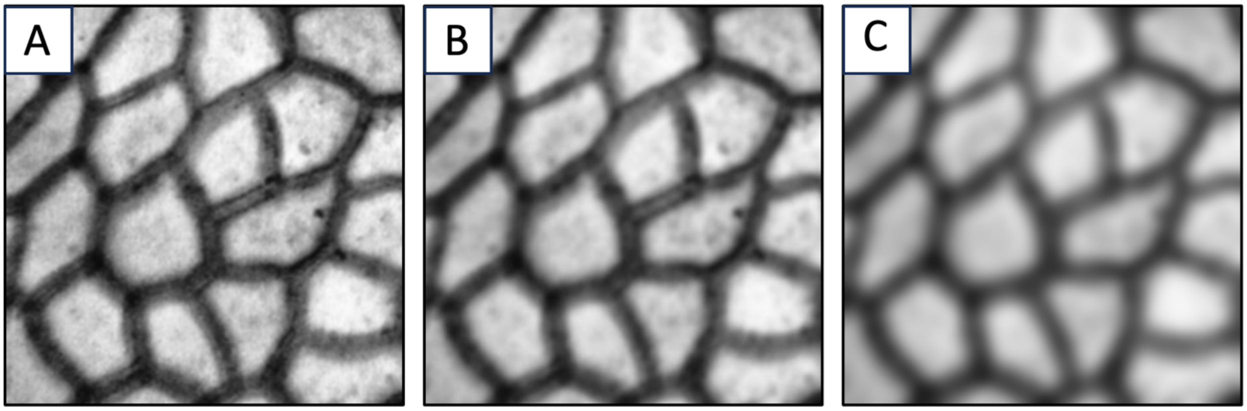

**Figure 20.** An image of *Castanea pumila* cuticle (FLMNH00115) after local contrast enhancement with gaussian blur set to **(A)** 1 pixel, **(B)** 2 pixels (optimal for this image), and **(C)** 5 pixels.

4. Apply Local Thresholding (Image > Adjust > Auto Local Threshold)
    a. In this step, you may experiment with different thresholding algorithms (Fig. 21) or select “Try All” for the method. This will result in a multi-paneled image showing the results of all thresholding algorithms for that image (Fig. 16). The “Try All” option is recommended for determining which thresholding methods work well on your dataset.

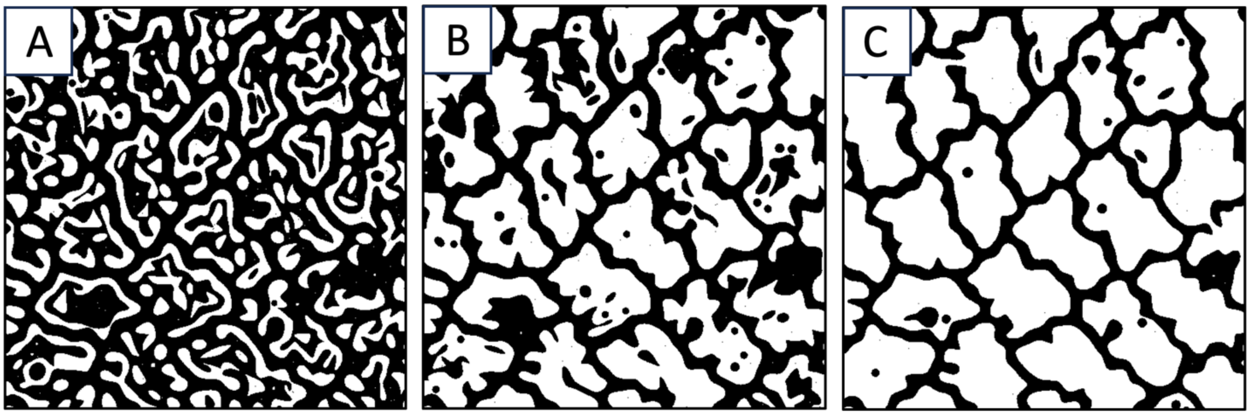

**Figure 21.** An image of *Talisia princeps* cuticle (FLMNH00443) after local contrast enhancement and blurring, thresholded by either the **(A)** NiBlack algorithm, **(B)** Bernsen algorithm, or **(C)** Sauvola algorithm (optimal for this image). See Lloyd et al. (2023) Fig. 2.

5. Apply shape smoothing (Plugins > Shape Smoothing)
    a. In this step, you may experiment with different smoothing values. The shape smoothing plugin from the BioMedGroup (Wagner, 2016) smoothes binary images by filtering a relative proportion of Fourier Descriptors of each "blob" within the image. More information can be found on the ImageJ website (https://imagej.net/plugins/shape-smoothing). An ideal smoothing value minimizes thresholding artifacts without obscuring gross cell shape (Fig. 22).

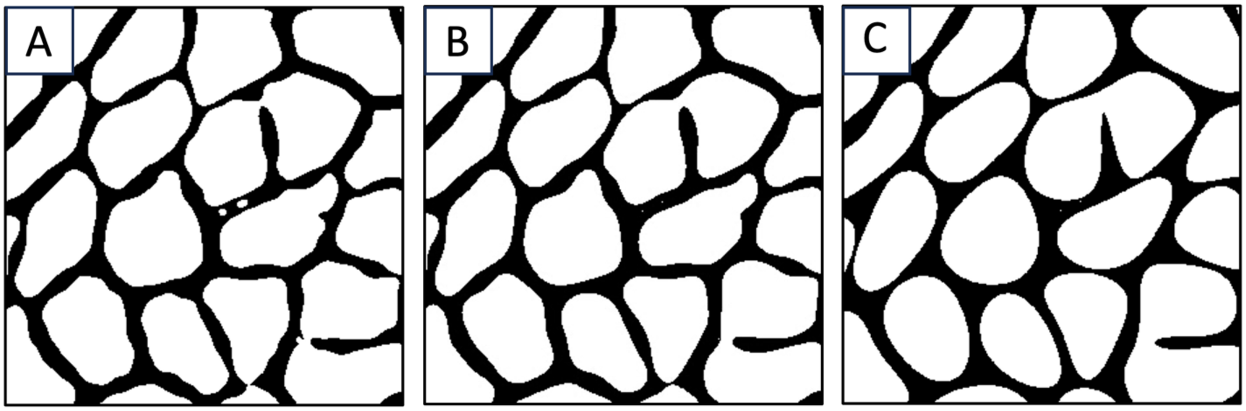

**Figure 22.** An image of *Castanea pumila* cuticle (FLMNH00115) after thresholding, with shape smoothing set to **(A)** 10% FDs retained, **(B)** 5% FDs retained (optimal for this image), and **(C)** 2% FDs retained.

6. As you experiment with these inputs, it is important to understand how a thresholded image will translate to a finalized ROI set.
    a. To take a thresholded image through the next steps performed by "Batch Generate ROIs," select the "Single Image Processor" macro (Plugins > Macros > CuticleTrace – Single Image Processor). By selecting

"Skeletonized Image (previously thresholded)" from the "Desired Outputs" menu, this macro generates a skeletonized image with 3-pixel-wide cell walls, which can be directly analyzed to generate ROIs (Fig. 23).

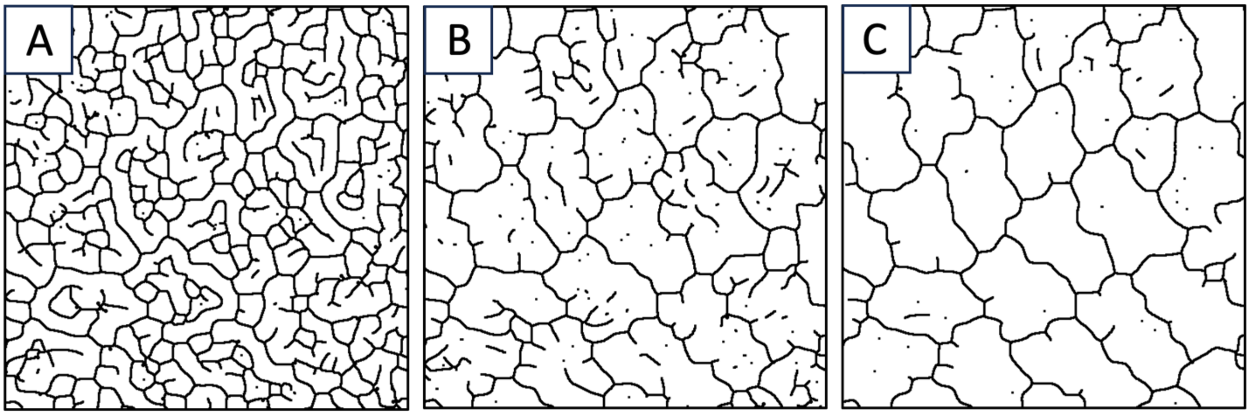

**Figure 23.** Skeletonized images of Talisia princeps cuticle (FLMNH00443) that has been thresholded by either the **(A)** NiBlack method, **(B)** Bernsen method, or **(C)** Sauvola method (optimal for this image). See Lloyd et al. (2023) Fig. 2.

b. To generate an unfiltered ROI set from a skeletonized image, continue to use the "Single Image Processor" macro (Plugins > Macros > CuticleTrace – Single Image Processor). By selecting "ROIs (previously skeletonized)" from the "Desired Outputs" menu, this macro generates an unfiltered refined and interpolated ROI set from a skeletonized image (Fig. 24).

Once you have decided upon input parameters by visually vetting ROI sets from a subset of images, you are ready to batch process images.

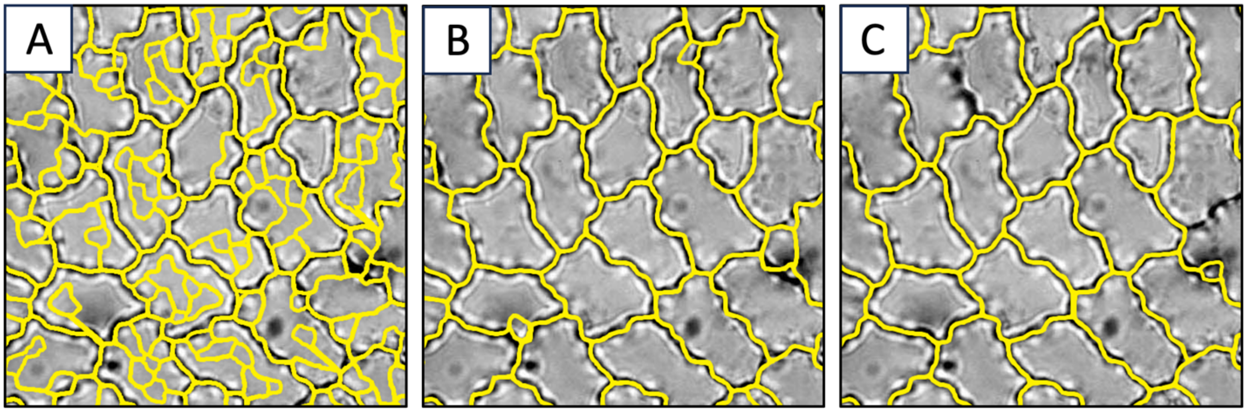

**Figure 24.** ROI sets generated from skeletonized images of Talisia princeps cuticle (FLMNH00443) that has been thresholded by either the **(A)** NiBlack method, **(B)** Bernsen method, or **(C)** Sauvola method (optimal for this image). See Lloyd et al. (2023) Fig. 2.

#### 4.2. Batch processing your own datasets

Now that you have an in-depth understanding of how to decide on your CuticleTrace input parameters, you are ready to apply CuticleTrace to your own images. Batch processing your own images is no different than batch processing the example dataset—just plug in your newly-determined input parameters and follow the tutorial from section 3.3.

With the CuticleTrace toolkit, we hope you generate increased sample sizes and more consistent measurements, all with significantly less time and effort in comparison to manual measurements. Good luck!
