## Supplementary material for "CuticleTrace: A toolkit for capturing cell outlines of leaf cuticle with implications for paleoecology and paleoclimatology": Figure S2

Supplemental Figure - Image plates. (left) Unfiltered; (center left) 2MADs; (center right) 1MAD; (right) Expert.

MLC_FLMNH00081_Fabaceae_Gleditsia_triacanthos_grp.jpg

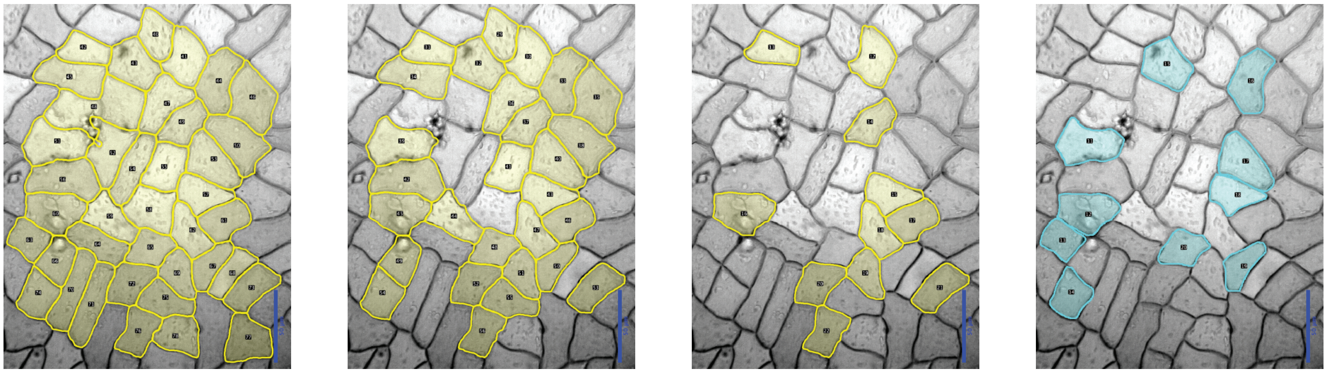

MLC_FLMNH00095_Fabaceae_Paubrasilia_echinata_grp_TaxUpd.jpg

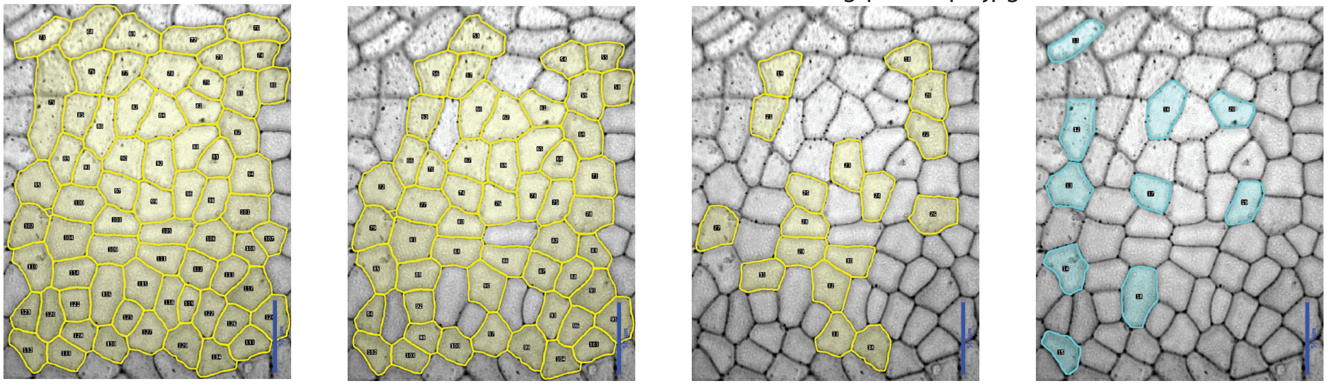

MLC_FLMNH00110_Fagaceae_Castanea_mollissima_grp.jpg

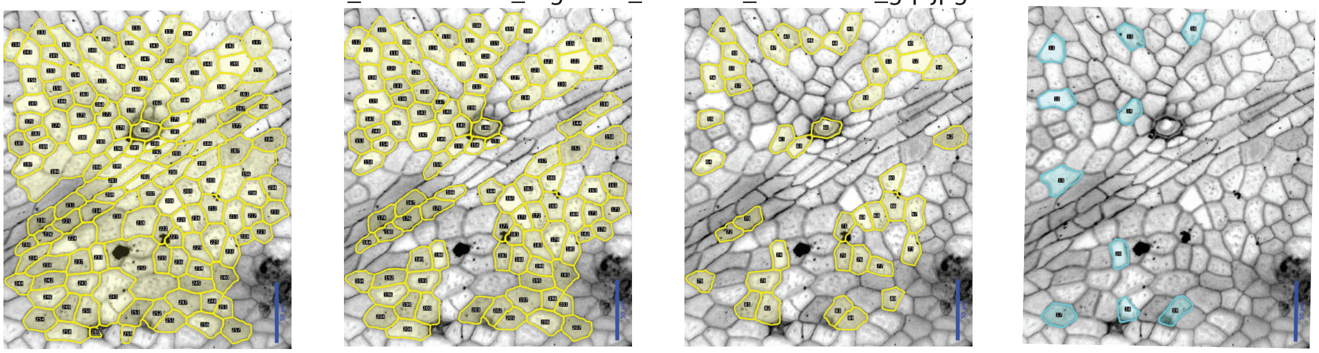

MLC_FLMNH00115_Fagaceae_Castanea_pumila_grp_TaxUpd.jpg

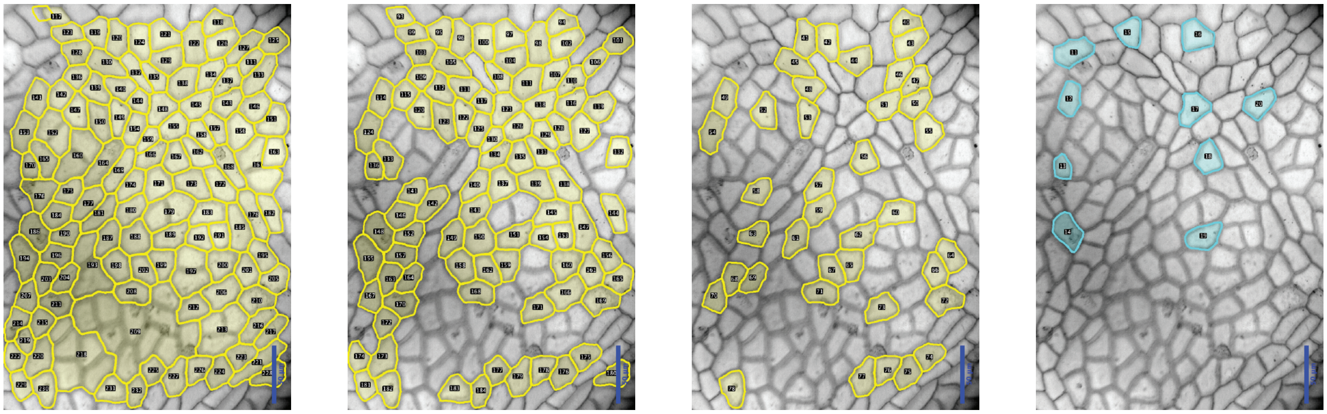

### Figure S2

Supplemental Figure - Image plates. (left) Unfiltered; (center left) 2MADs; (center right) 1MAD; (right) Expert.

MLC_FLMNH00146_Fagaceae_Quercus_sp_grp_TaxUpd.jpg

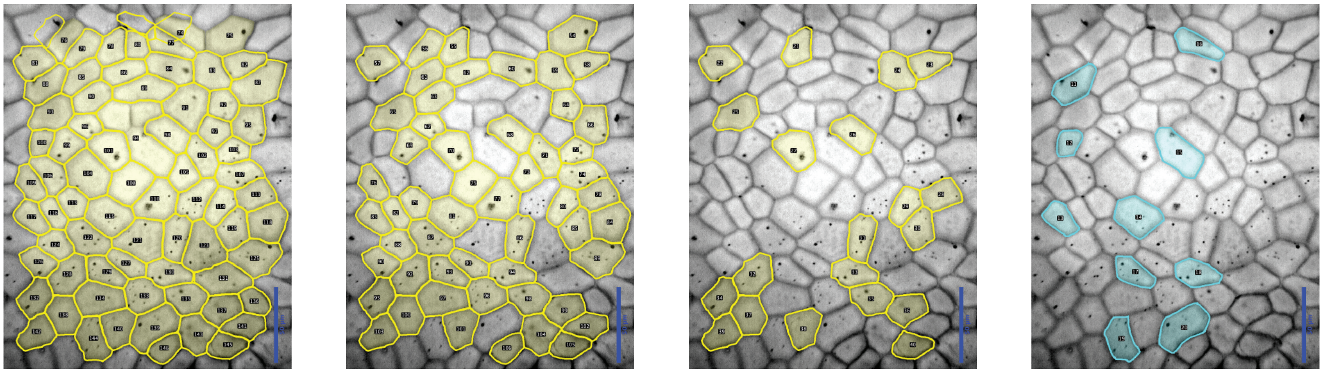

MLC_FLMNH00170_Lauraceae_Beilschmiedia_costaricensis_grp.jpg

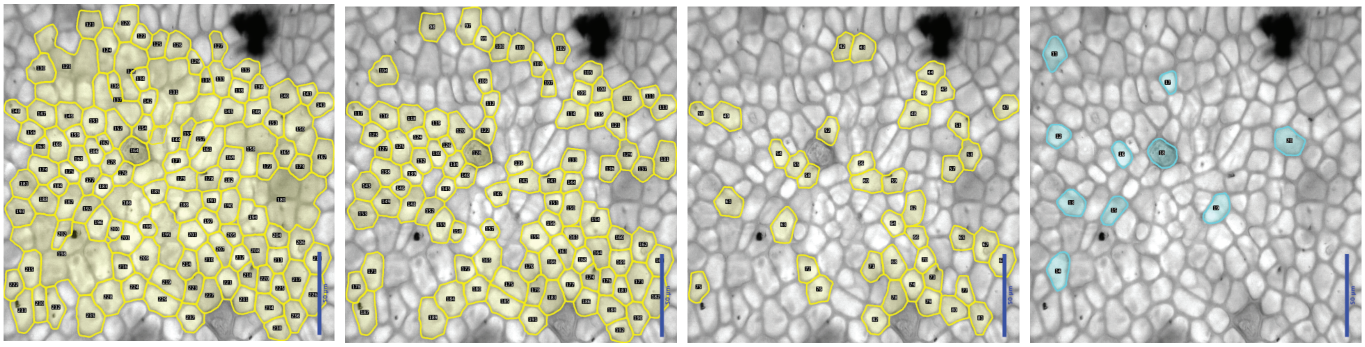

MLC_FLMNH00174_Lauraceae_Aniba_burchelli_gray_400dpi_TaxUpd.jpg

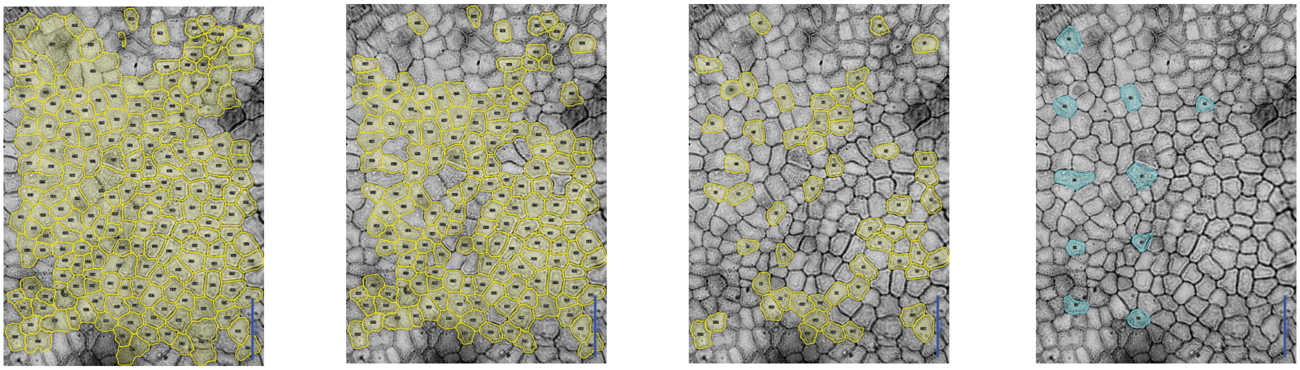

MLC_FLMNH00175_Lauraceae_Aiouea_impressa_grouped_gray_400dpi.jpg

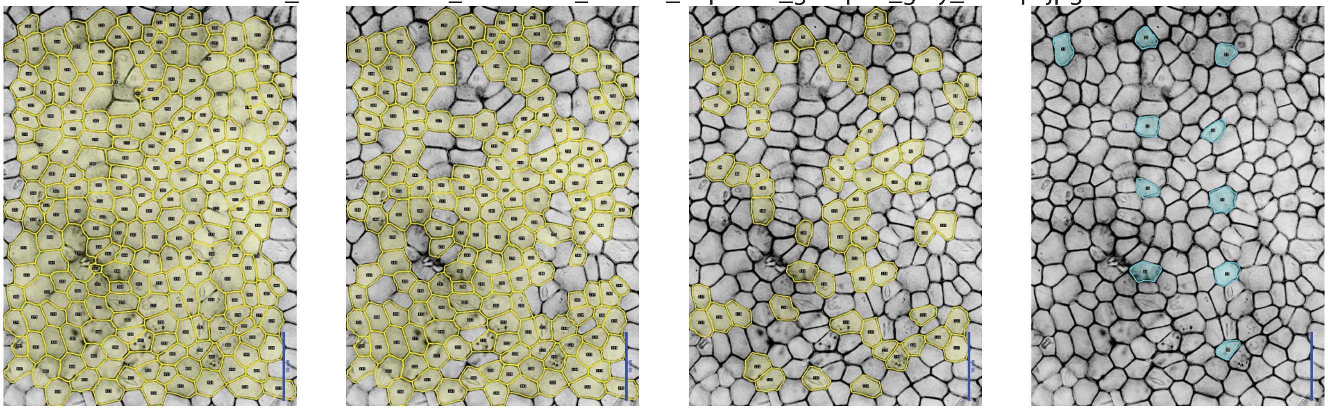

### Figure S2

Supplemental Figure - Image plates. (left) Unfiltered; (center left) 2MADs; (center right) 1MAD; (right) Expert.

MLC_FLMNH00177_Lauraceae_Ocotea_insularis_gray_400dpi_TaxUpd.jpg

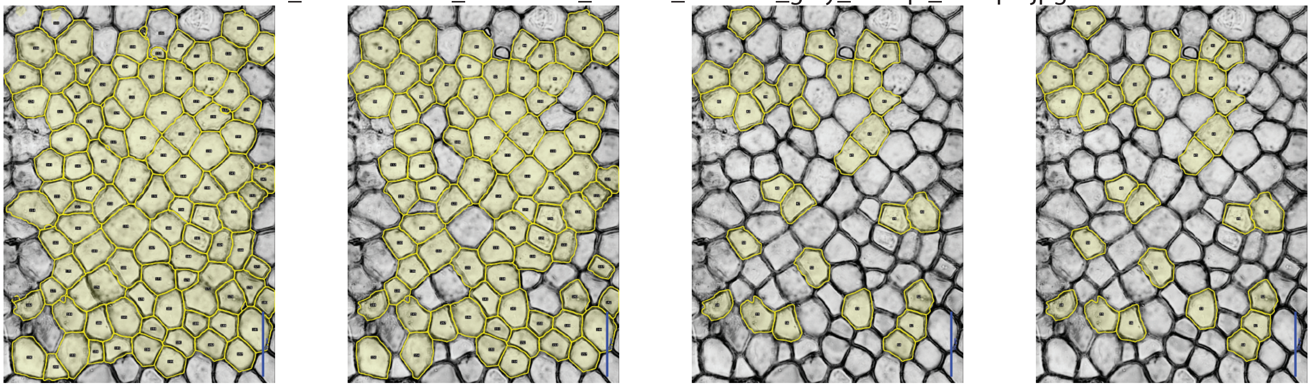

MLC_FLMNH00178_Lauraceae_Damburneya_salicifolia_gray_400dpi_TaxUpd.jpg

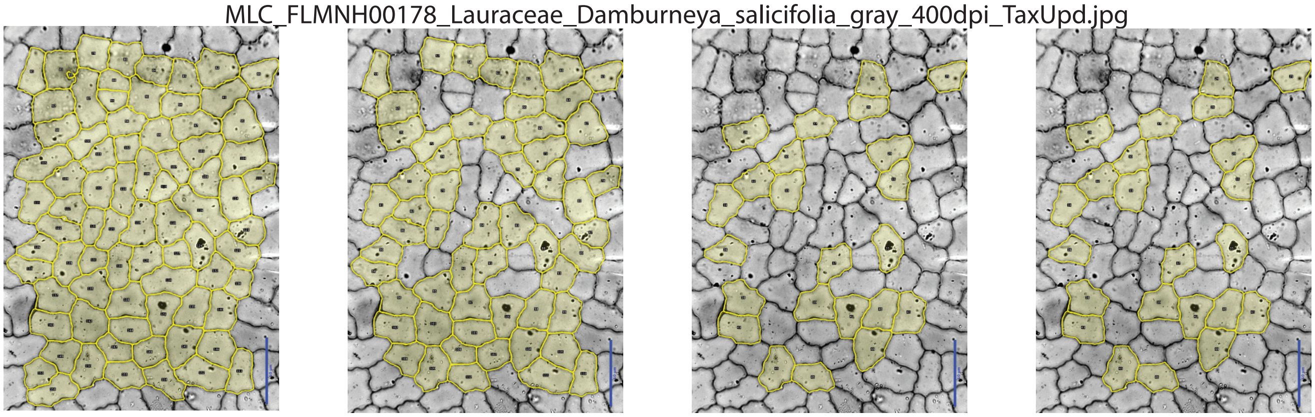

MLC_FLMNH00183_Lauraceae_Ocotea_tenuiflora_grouped_gray_400dpi.jpg

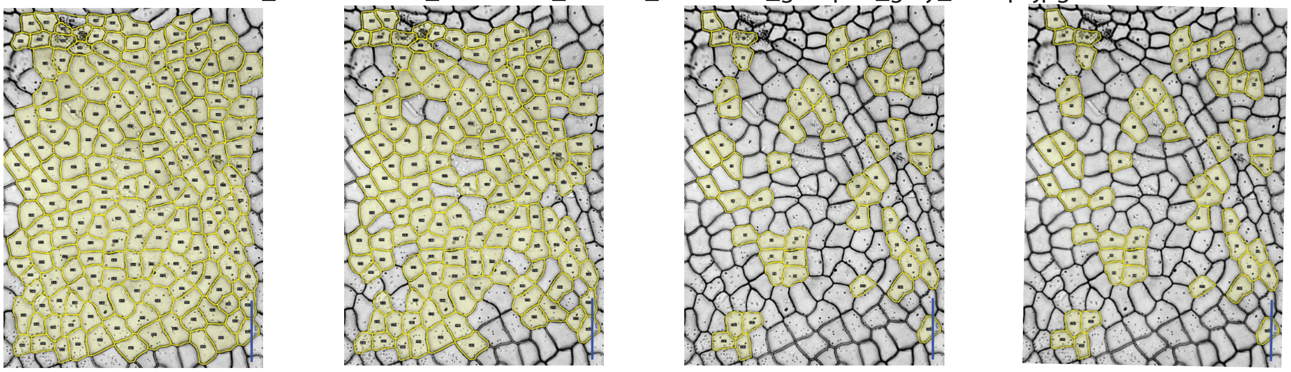

MLC_FLMNH00185_Lauraceae_Ocotea_tarapotana_grouped_gray_400dpi.jpg

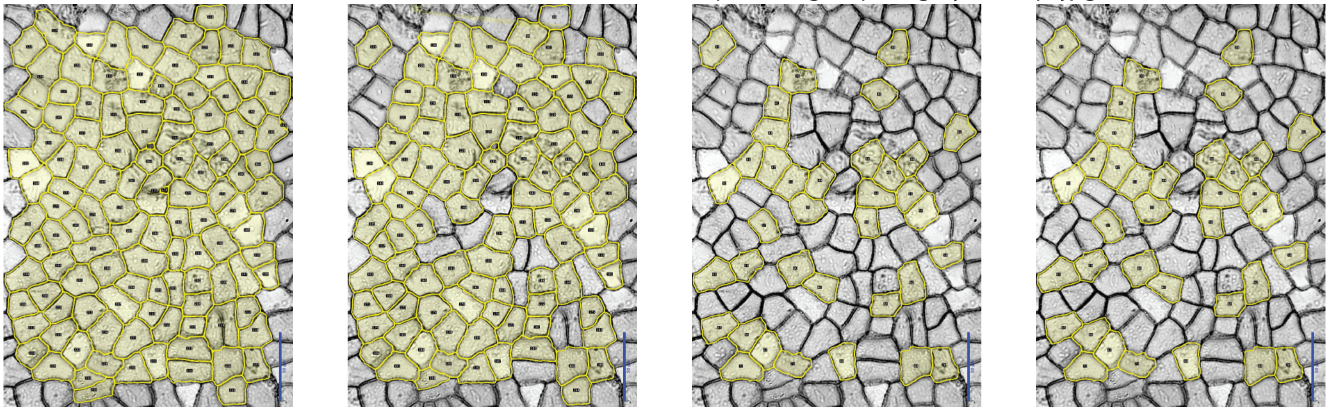

#### Figure S2

Supplemental Figure - Image plates. (left) Unfiltered; (center left) 2MADs; (center right) 1MAD; (right) Expert.

MLC_FLMNH00186_Lauraceae_Ocotea_suaveolens_grouped_gray_400dpi_TaxUpd.jpg

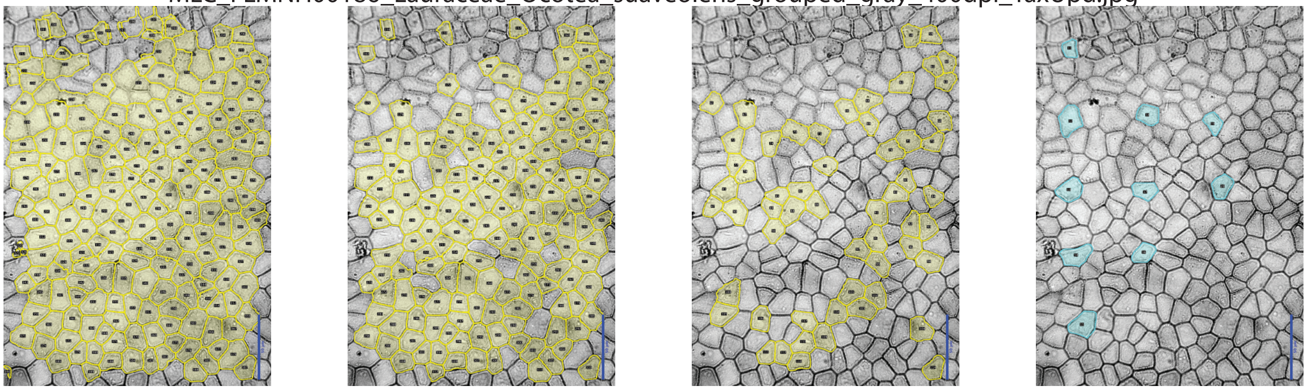

MLC_FLMNH00235_Lauraceae_Ocotea_spathulata_grouped_gray_400dpi.jpg

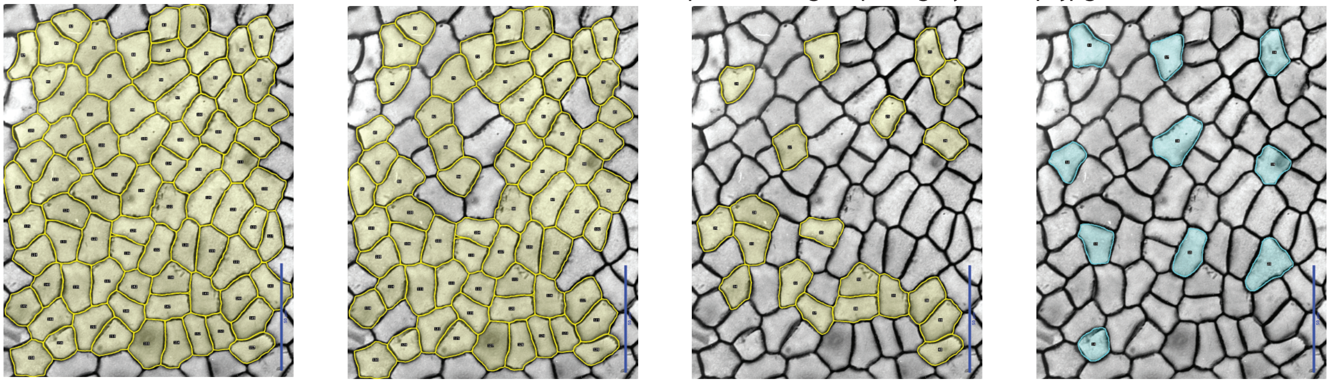

MLC_FLMNH00260_Lauraceae_Nectandra_oppositifolia_gray_400dpi_TaxUpd.jpg

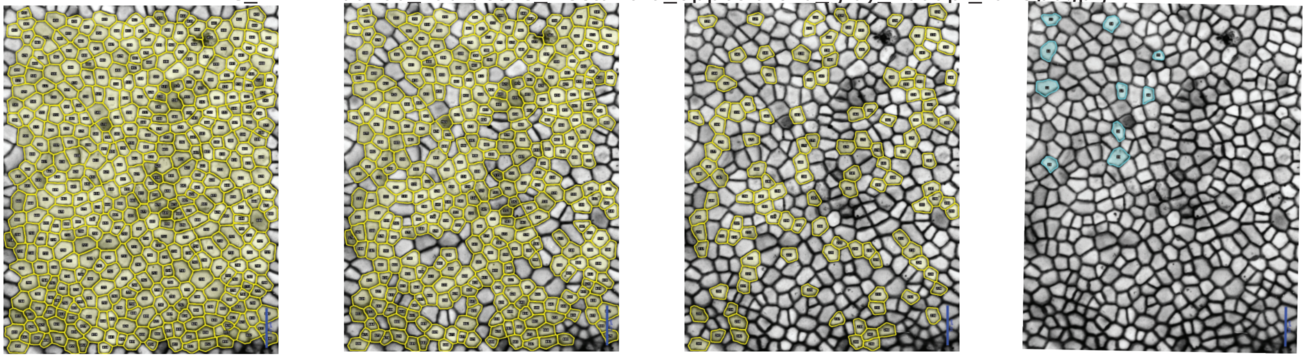

MLC_FLMNH00309_Sapindaceae_Cubilia_cubili_grp_TaxUpd.jpg

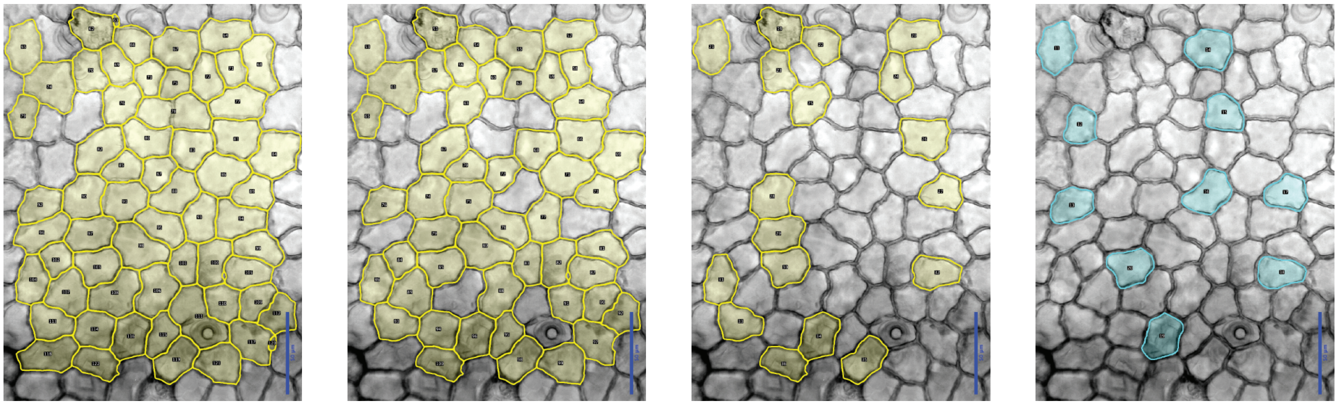

### Figure S2

Supplemental Figure - Image plates. (left) Unfiltered; (center left) 2MADs; (center right) 1MAD; (right) Expert.

MLC_FLMNH00310_Sapindaceae_Cupania_glabra_grp.jpg

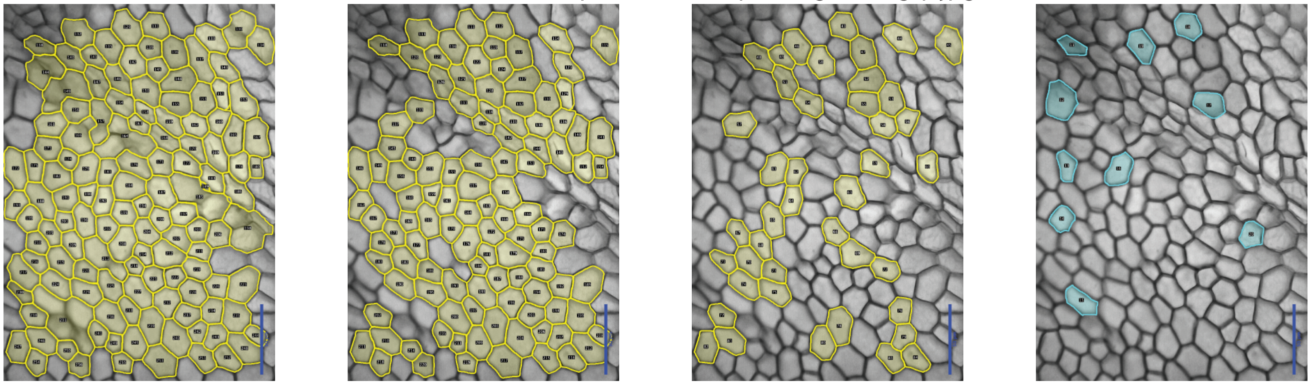

MLC_FLMNH00315_Sapindaceae_Elattostachys_apetala_grp_TaxUpd.jpg

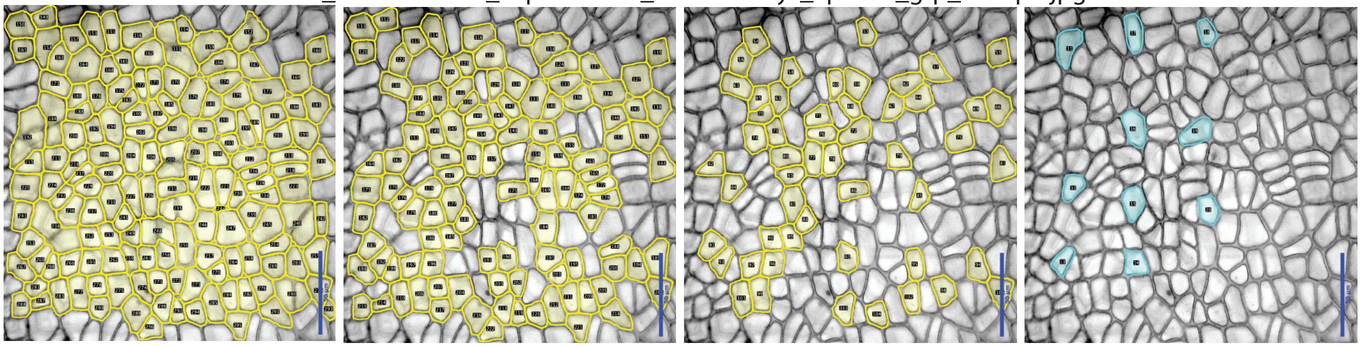

MLC_FLMNH00377_Sapindaceae_Thouinidium_pinnatum_grouped_gray_400dpi.jpg

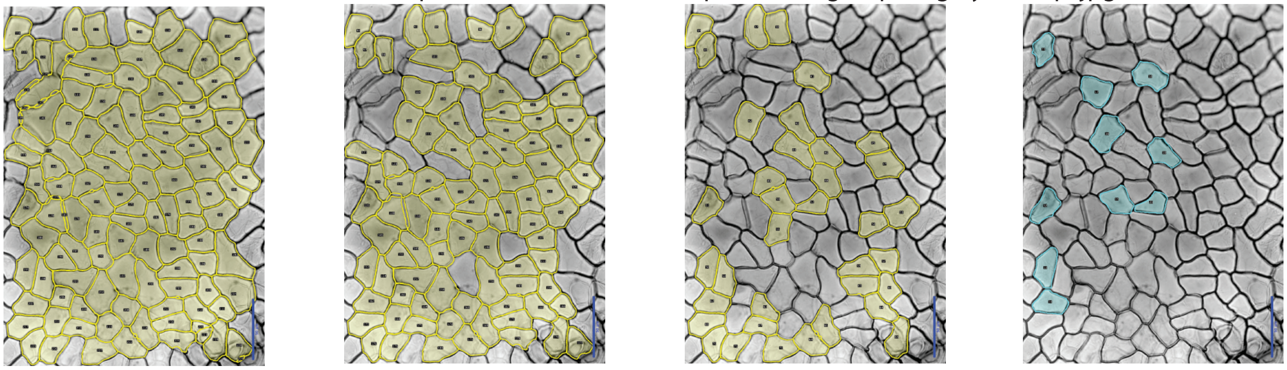

MLC_FLMNH00382_Sapindaceae_Toulicia_laevigata_grouped_gray_400dpi.jpg

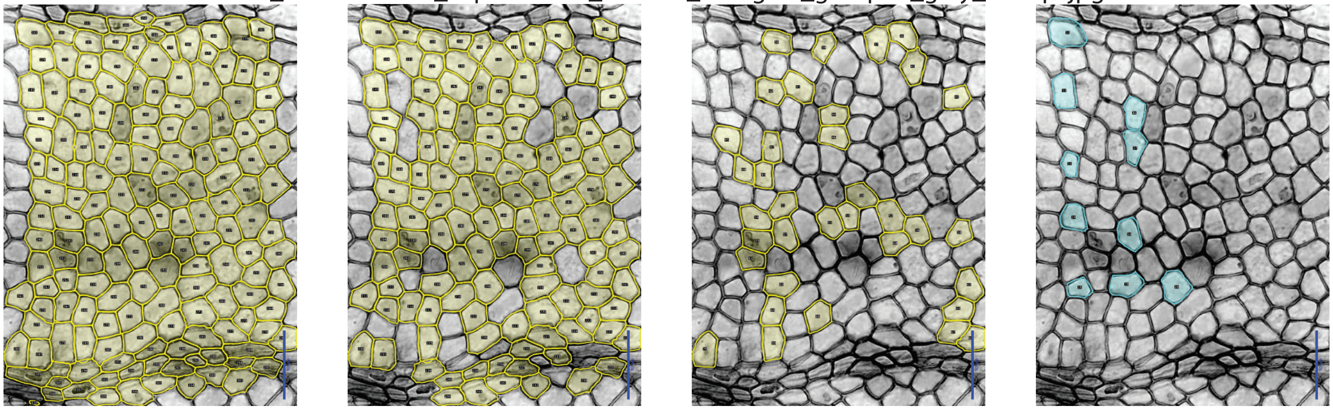

### Figure S2

Supplemental Figure - Image plates. (left) Unfiltered; (center left) 2MADs; (center right) 1MAD; (right) Expert.

MLC_FLMNH00411_Sapindaceae_Cupania_rigida_grouped_gray_400dpi.jpg

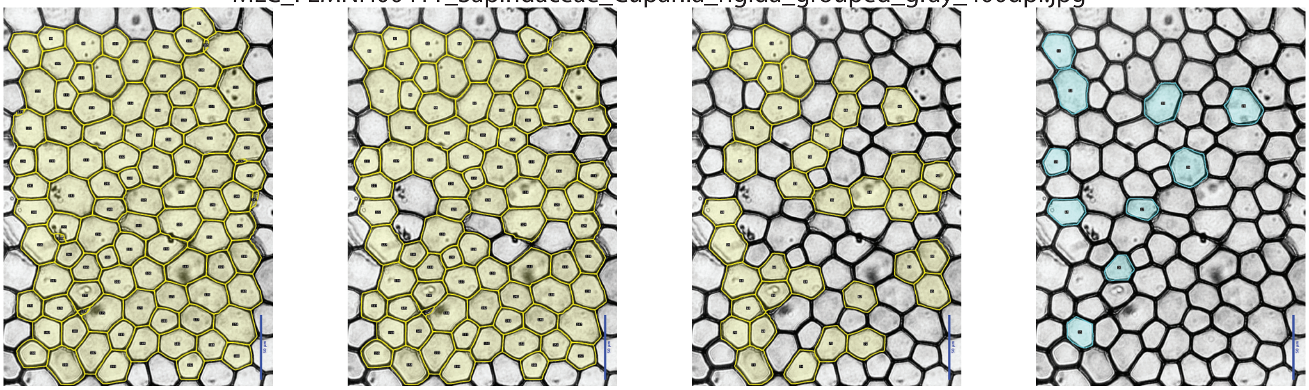

MLC_FLMNH00424_Sapindaceae_Cupania_glabra_grouped_gray_400dpi_TaxUpd.jpg

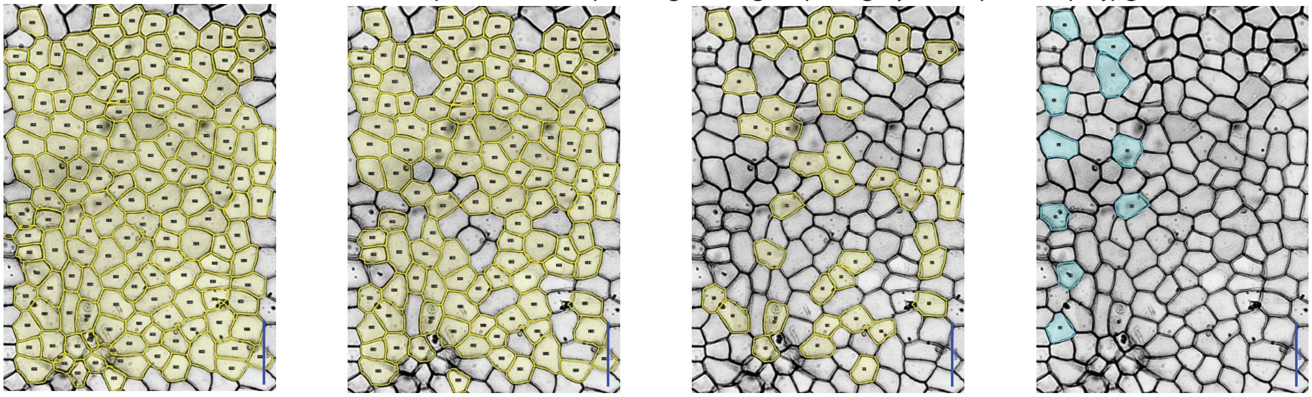

MLC_FLMNH00443_Sapindaceae_Talisia_princeps_grouped_gray_400dpi_TaxUpd.jpg

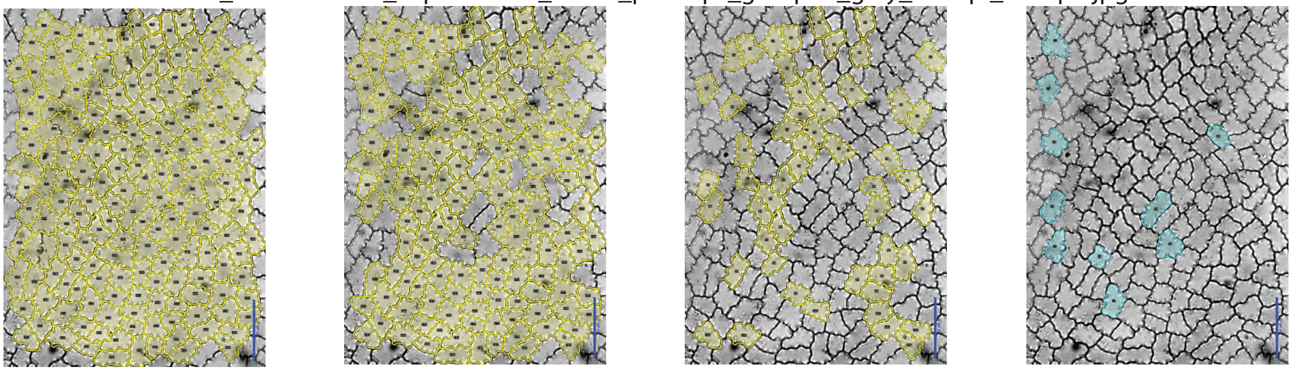

MLC_FLMNH00444_Sapindaceae_Talisia_esculenta_grouped_gray_400dpi.jpg

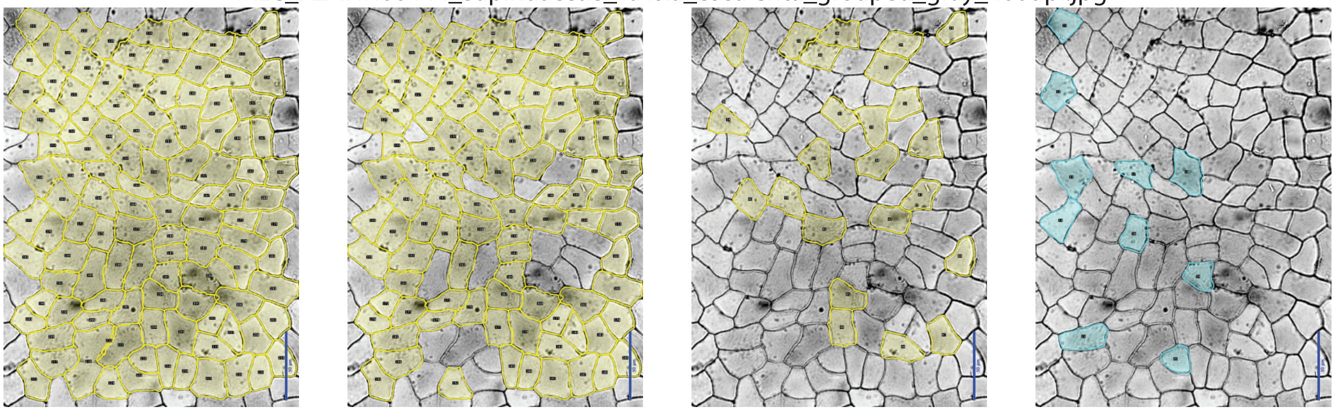

### Figure S2

Supplemental Figure - Image plates. (left) Unfiltered; (center left) 2MADs; (center right) 1MAD; (right) Expert.

MLC_FLMNH00458_Meliaceae_Cedrela_odorata_grp.jpg

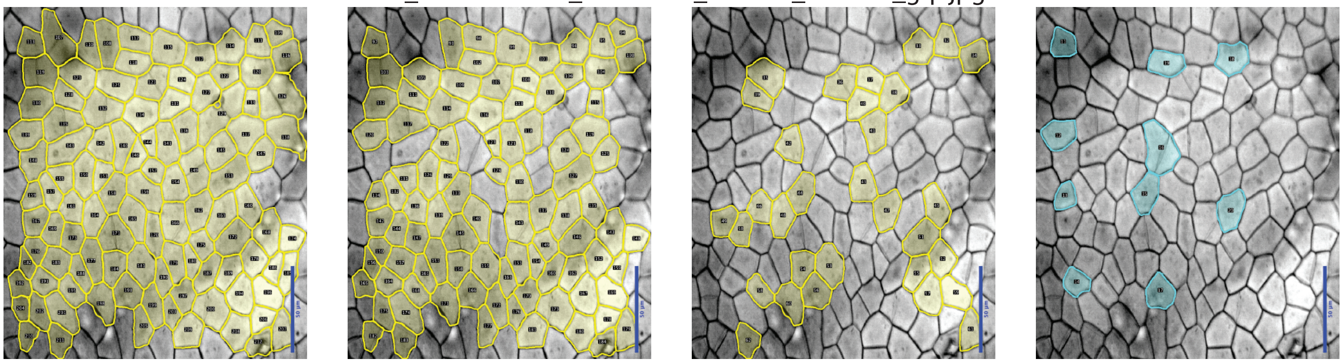

MLC_FLMNH00510_Anacardiaceae_Toxicodendron_striatum_grp_TaxUpd.jpg

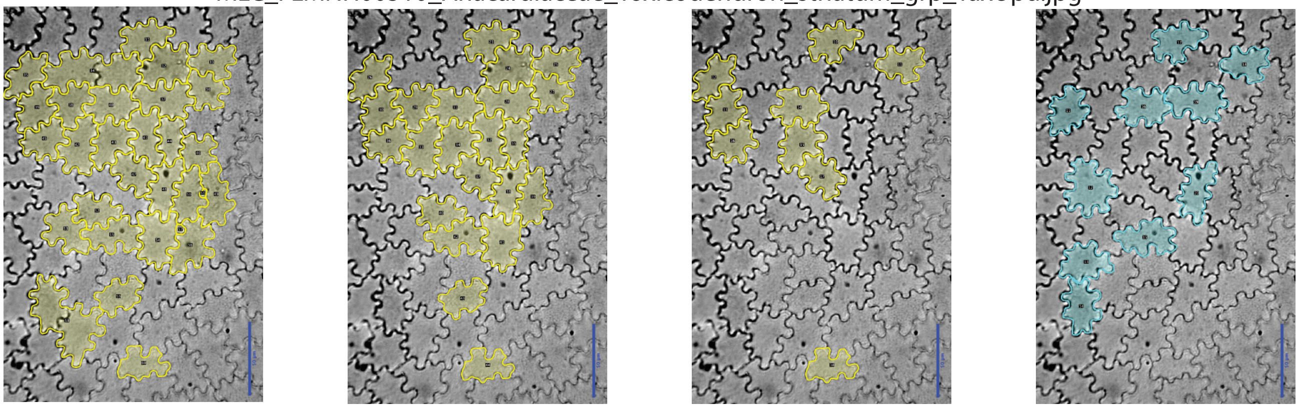

MLC_FLMNH00520_Rhamnaceae_Frangula_capreifolia_grp_TaxUpd.jpg

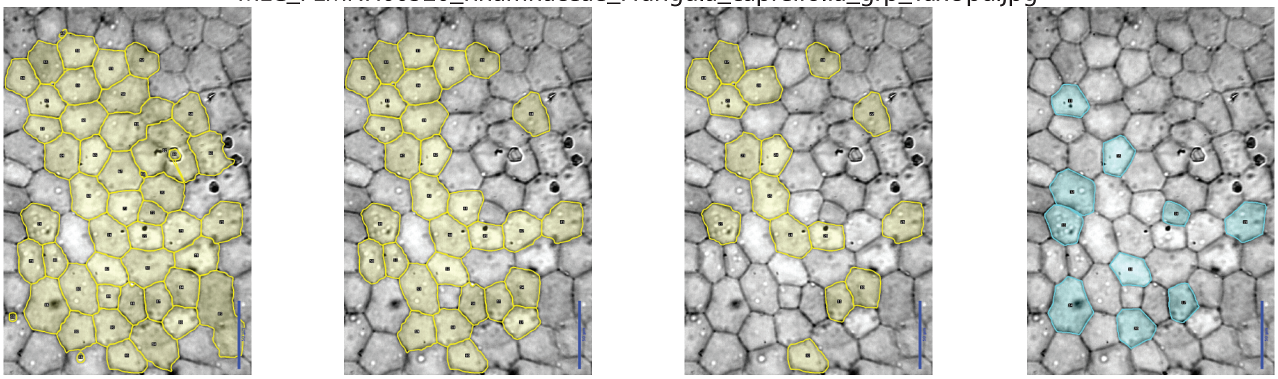

MLC_FLMNH00551_Fabaceae_Gymnocladus_dioica_grp.jpg

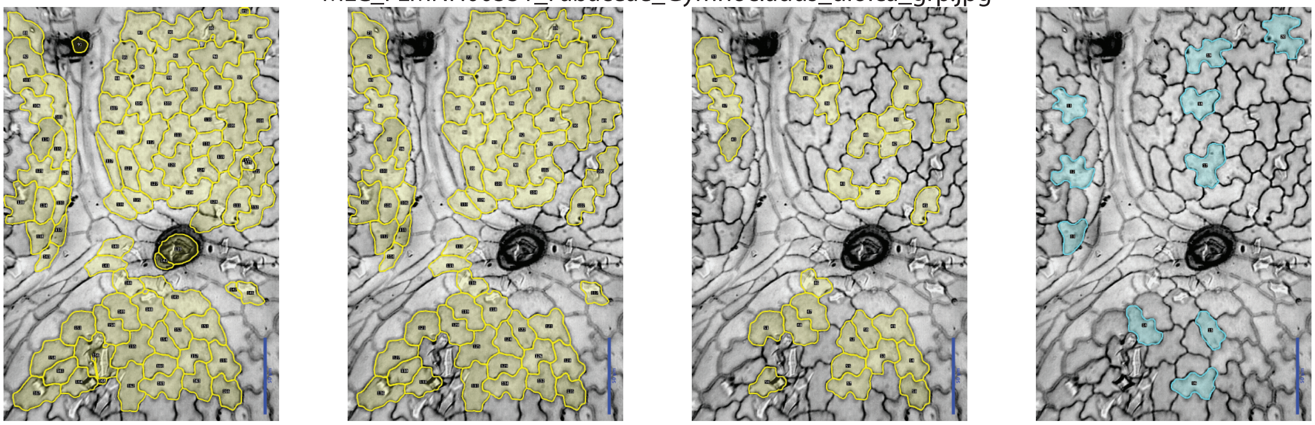

### Figure S2

Supplemental Figure - Image plates.  (left) Unfiltered; (center left) 2MADs; (center right) 1MAD; (right) Expert.

MLC_FLMNH00561_Fabaceae_Gleditsia_triacanthos_grp.jpg

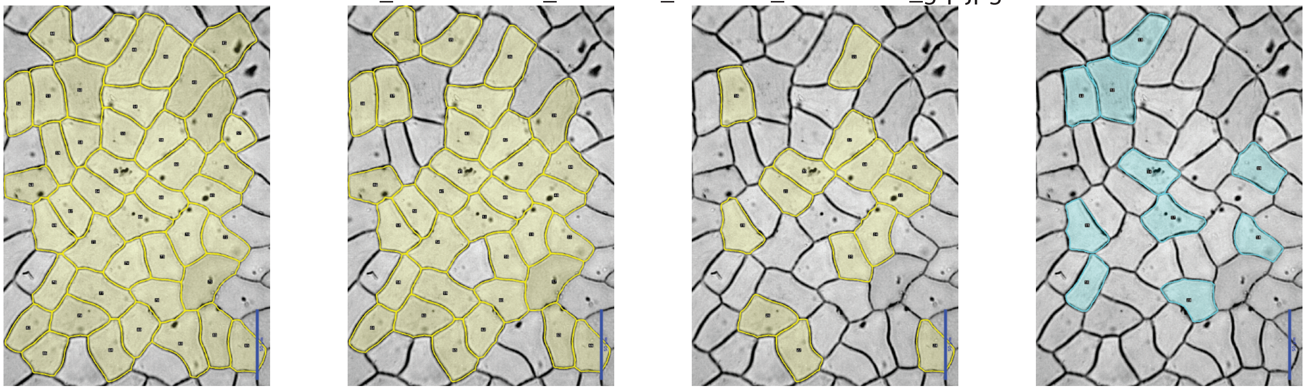

MLC_FLMNH00587_Araceae_Philodendron_sodiroi_grp.jpg

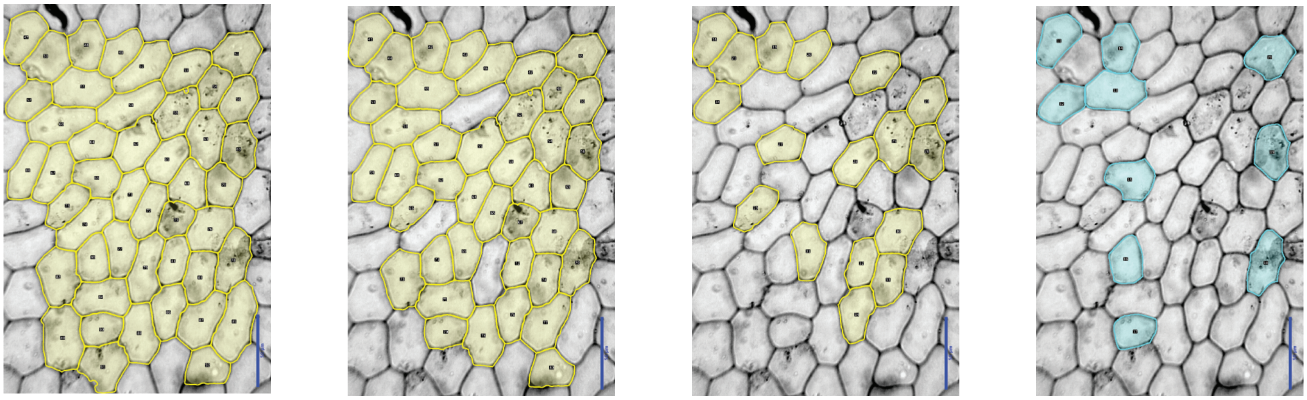

MLC_FLMNH00616_Fagaceae_Quercus_dumosa_grp.jpg

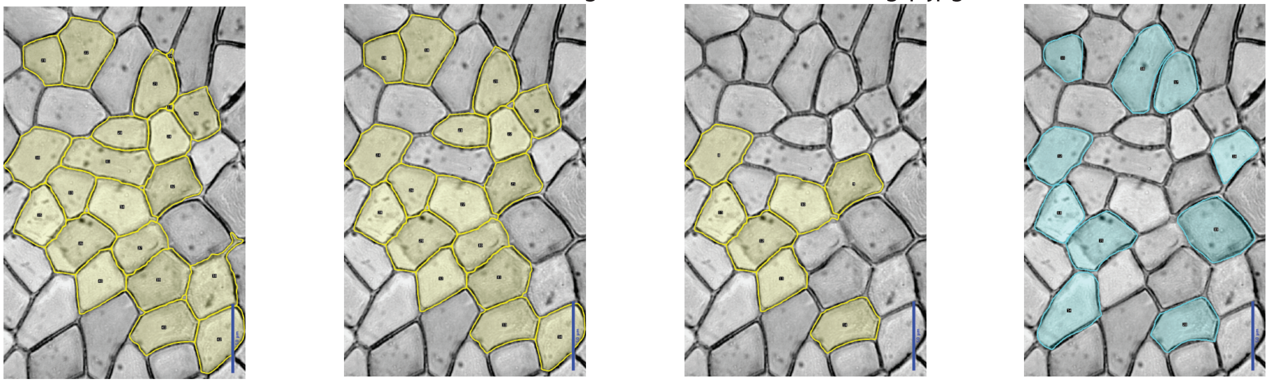

MLC_FLMNH00626_Fagaceae_Quercus_affinis_grp.jpg

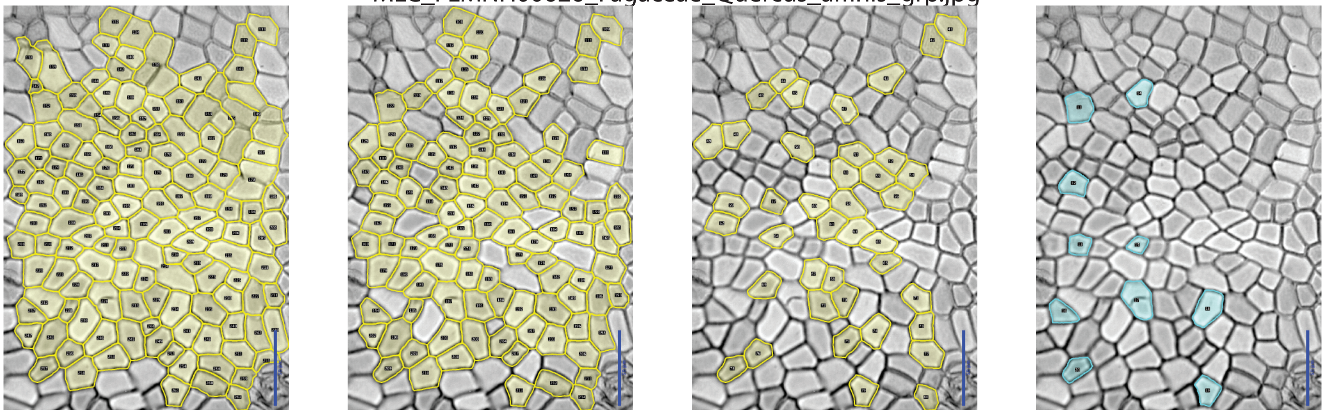

### Figure S2

Supplemental Figure - Image plates. (left) Unfiltered; (center left) 2MADs; (center right) 1MAD; (right) Expert.

MLC_FLMNH00652_Araceae_Chamaedorea_brachypoda_grp.jpg

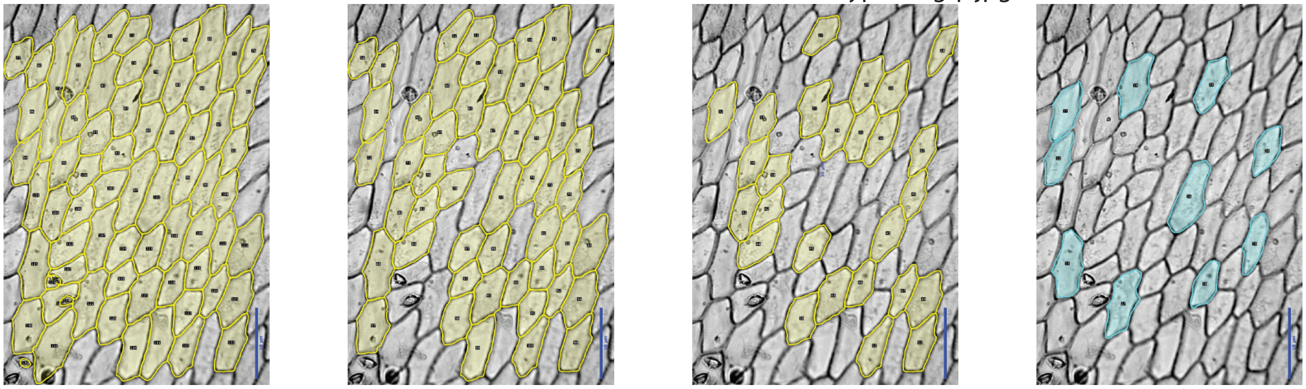

MLC_FLMNH00711_Araceae_Xanthosoma_sagittifolium_grp_TaxUpd.jpg

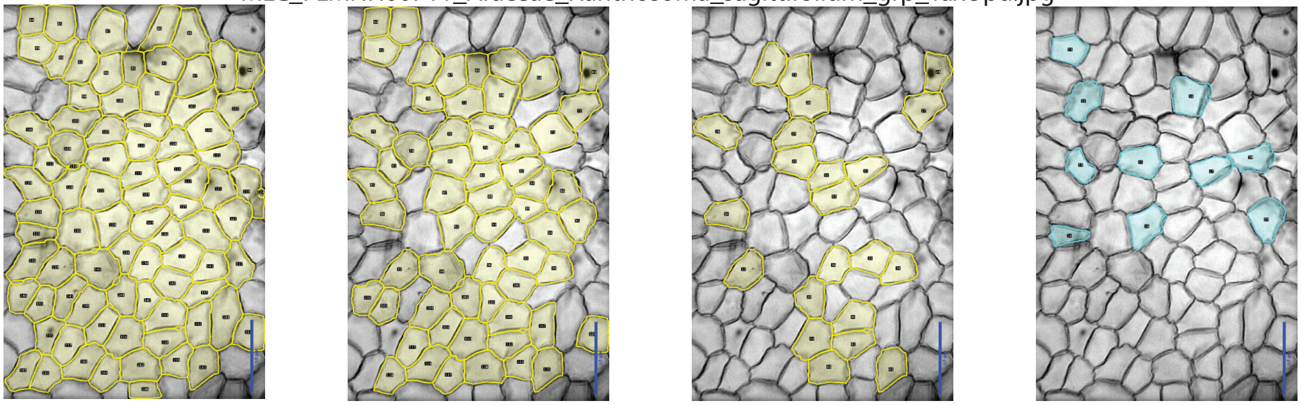

MLC_FLMNH00714_Aceraceae_Acer_skutchii_grp.jpg

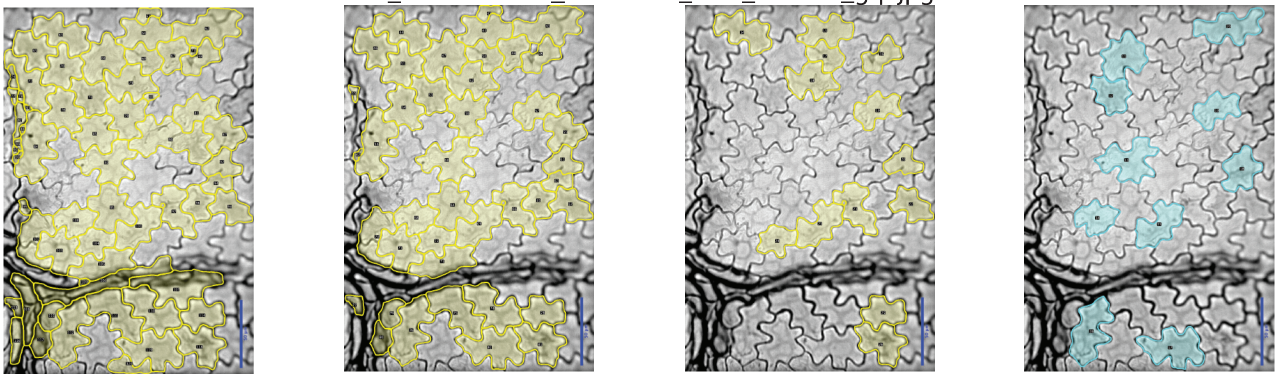

MLC_FLMNH00716_Myricaceae_Myrica_cerifera_grp_TaxUpd.jpg

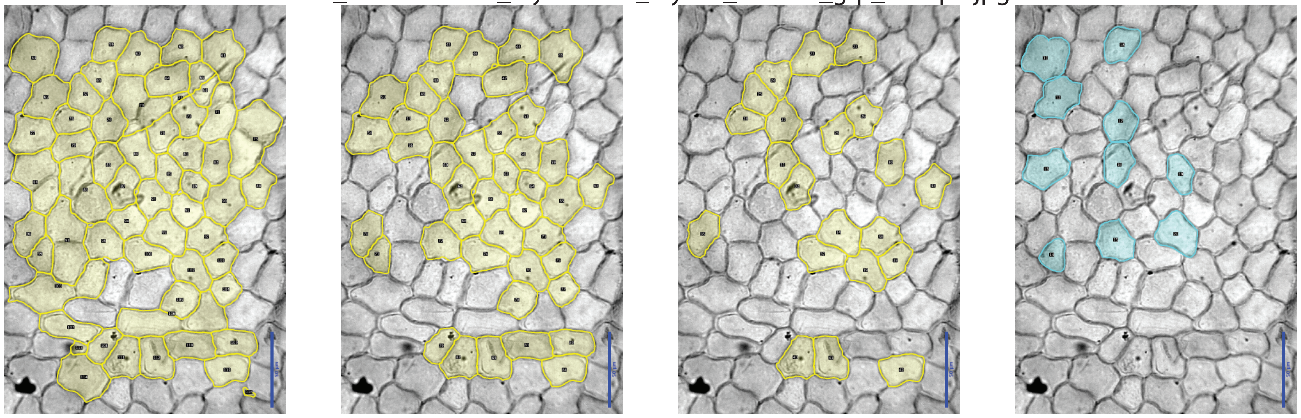

### Figure S2

Supplemental Figure - Image plates. (left) Unfiltered; (center left) 2MADs; (center right) 1MAD; (right) Expert.

MLC_FLMNH00767_Araliaceae_Oreopanax_capitatus_grp_TaxUpd.jpg

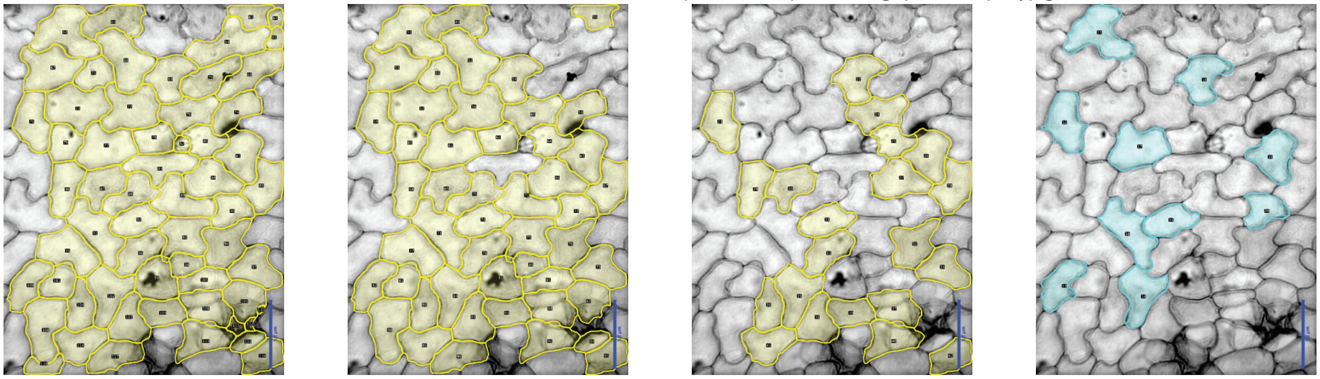

MLC_FLMNH00783_Proteaceae_Knightia_excelsa_grp.jpg

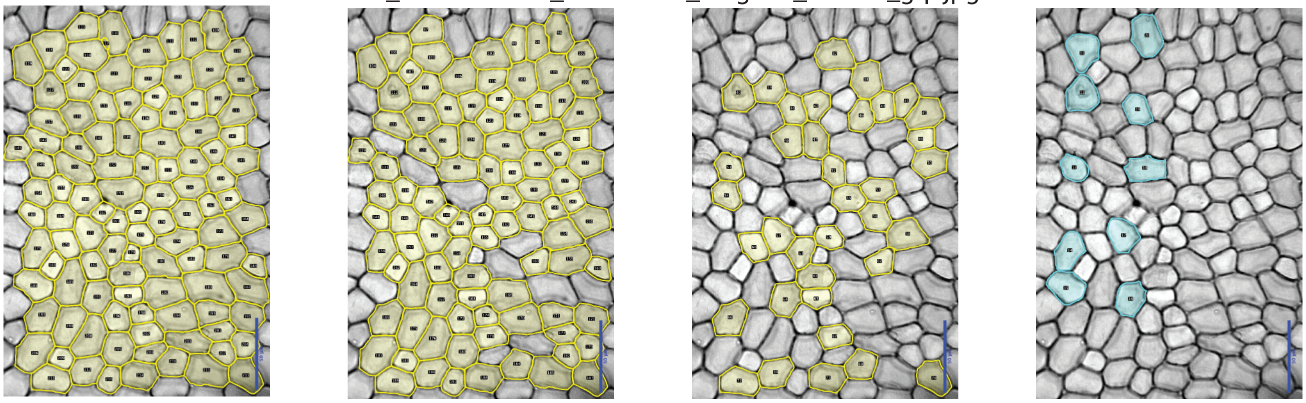

MLC_FLMNH00787_Monimiaceae_Hedycarya_arborea_grp.jpg

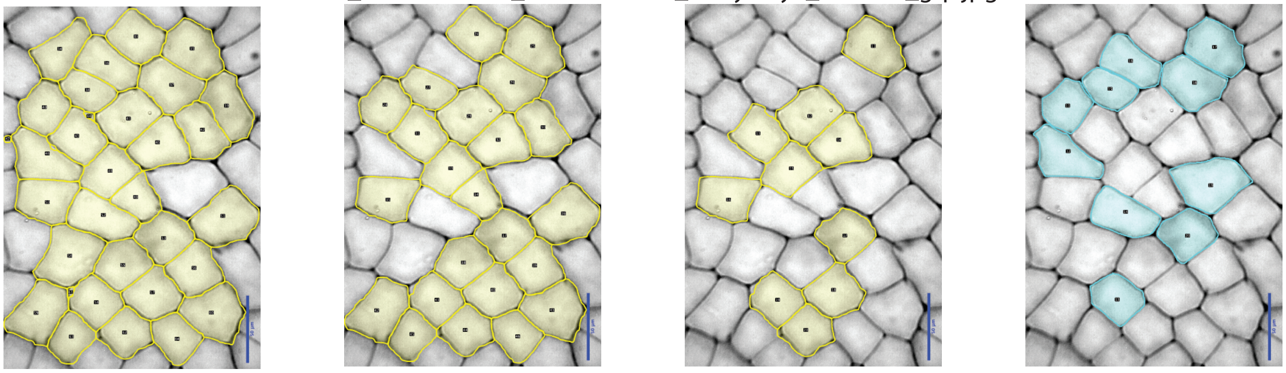

MLC_FLMNH00844_Picramniaceae_Picramnia_antidesma_subsp_fessonia_grp_TaxUpd.jpg

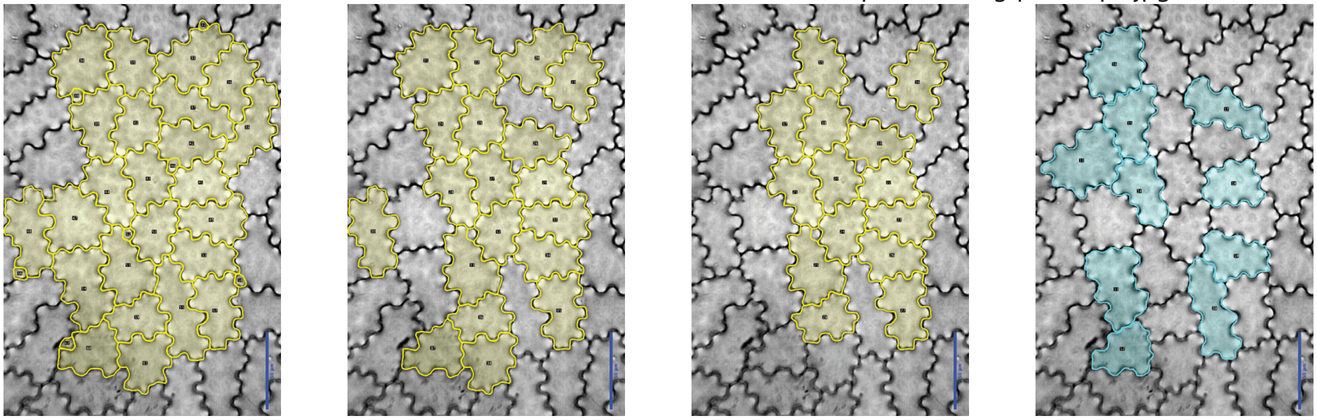

#### Figure S2

Supplemental Figure - Image plates. (left) Unfiltered; (center left) 2MADs; (center right) 1MAD; (right) Expert.

MLC_FLMNH00845_Picramniaceae_Picramnia_latifolia_grp_TaxUpd.jpg

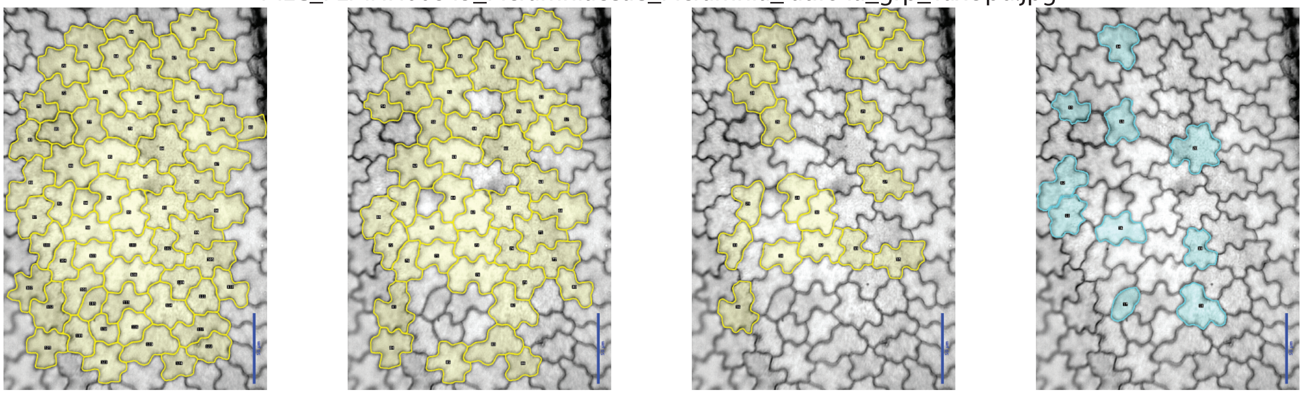

MLC_FLMNH00850_Meliaceae_Guarea_bijuga_grp.jpg

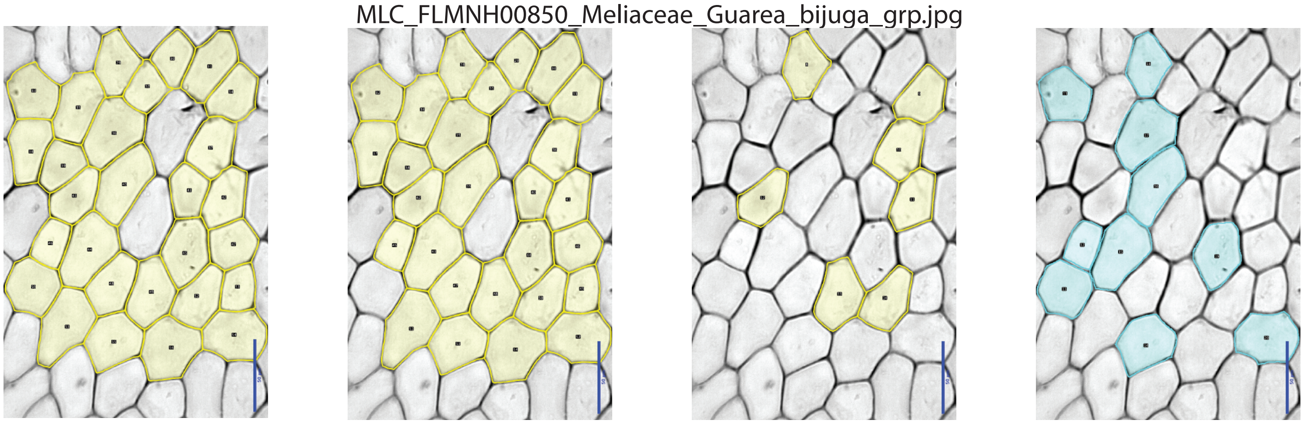

MLC_FLMNH00997_Primulaceae_Ardisia_bracteosa_grouped_gray_400dpi_TaxUpd.jpg.jpg

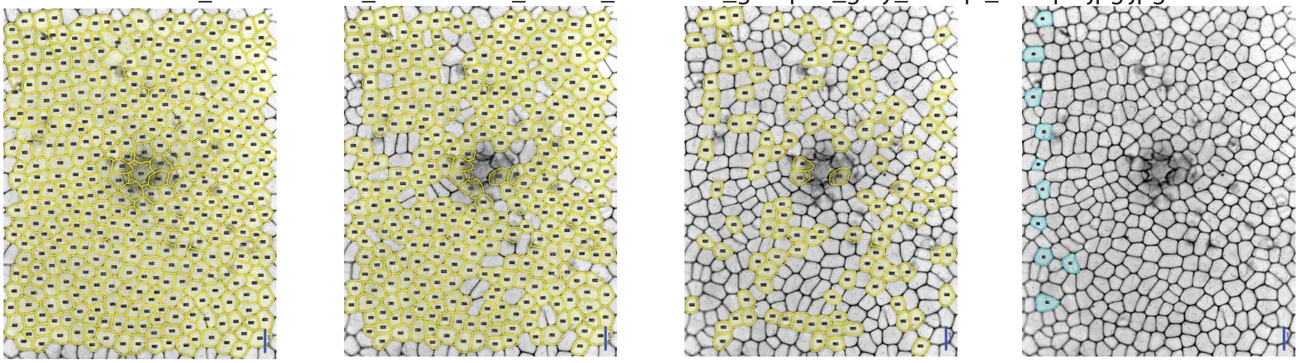

MLC_FLMNH01844_Annonaceae_Guatteria_dolichopoda_grp_TaxUpd.jpg

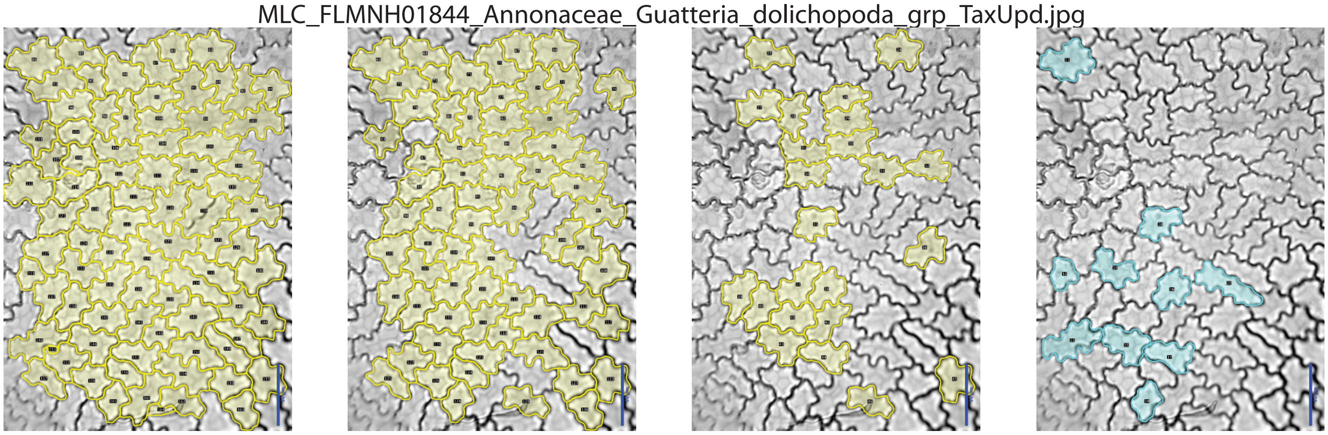

### Figure S2

Supplemental Figure - Image plates. (left) Unfiltered; (center left) 2MADs; (center right) 1MAD; (right) Expert.

MLC_FLMNH02064_Myristicaceae_Myristica_agusanensis_grp.jpg

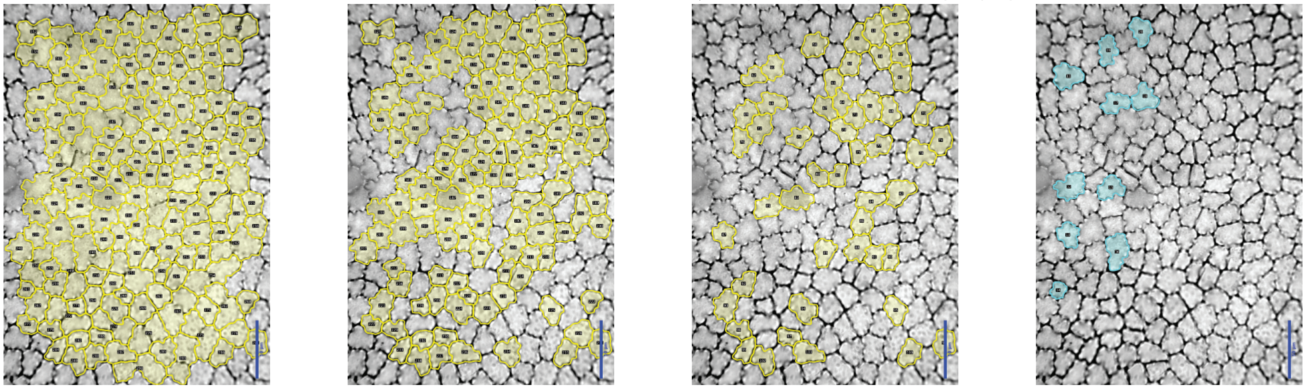

MLC_FLMNH02269_Apocynaceae_Aspidosperma_megalocarpon_grp.jpg

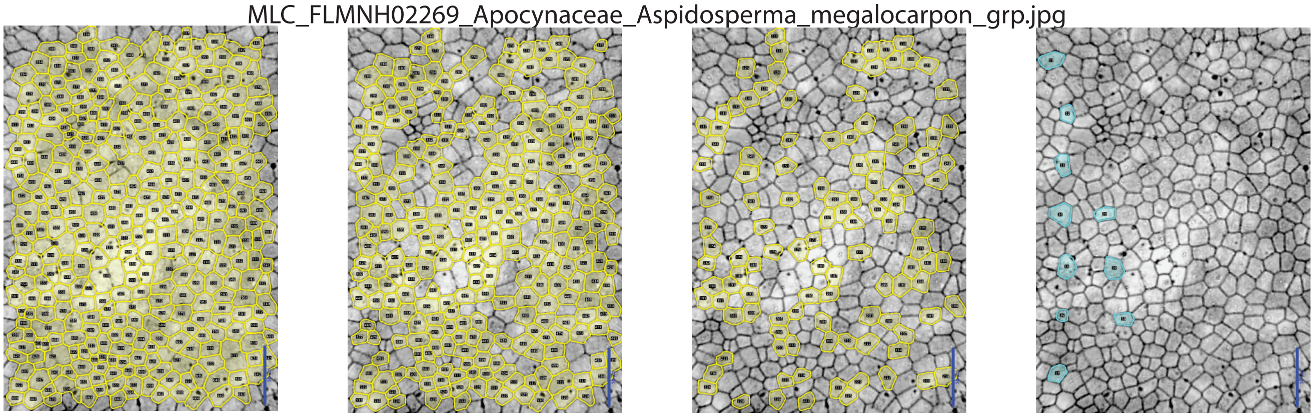

MLC_FLMNH02450_Rubiaceae_Condaminea_corymbosa_grp.jpg

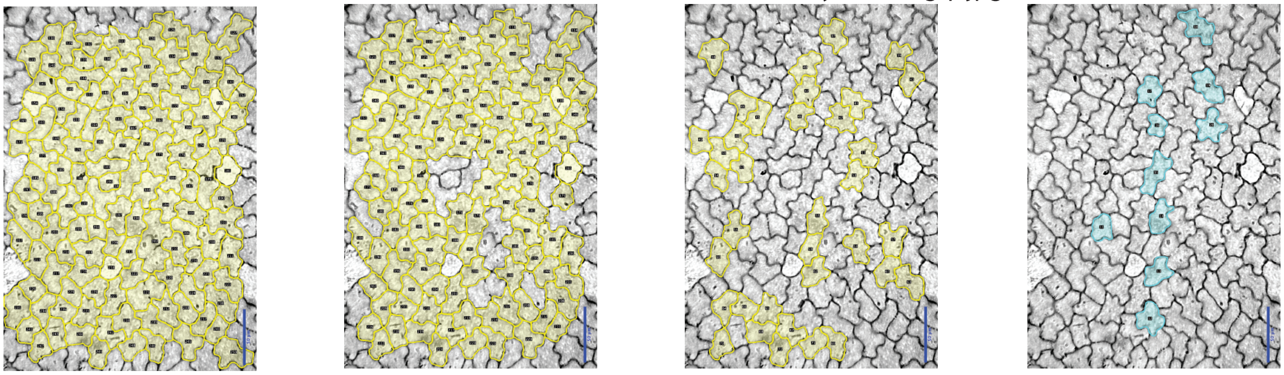

MLC_FLMNH02589_Annonaceae_Anaxagorea_petiolata_grp.jpg

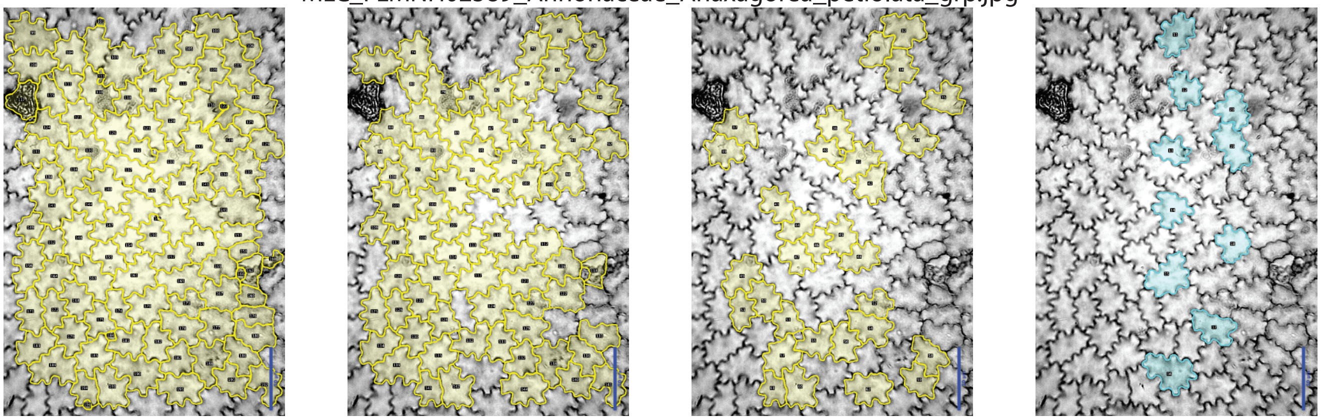

#### Figure S2

Supplemental Figure - Image plates.  (left) Unfiltered; (center left) 2MADs; (center right) 1MAD; (right) Expert.

MLC_FLMNH05215_Sapotaceae_Pouteria_durlandii_grp.jpg

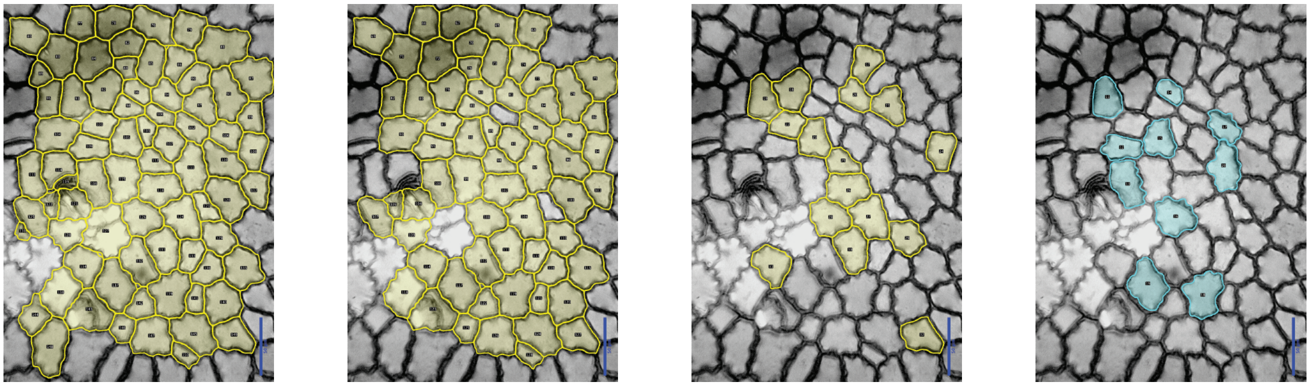

MLC_FLMNH05326_Sapindaceae_Cupania_livida_grp.jpg

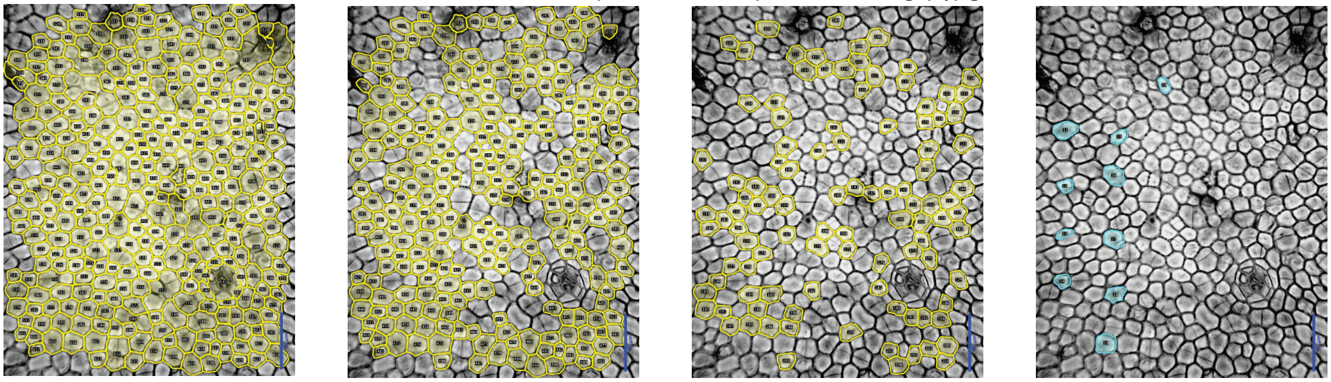
